# Deciphering 3’ UTR mediated gene regulation using interpretable deep representation learning

**DOI:** 10.1101/2023.09.08.556883

**Authors:** Yuning Yang, Gen Li, Kuan Pang, Wuxinhao Cao, Xiangtao Li, Zhaolei Zhang

## Abstract

The 3’untranslated regions (3’UTRs) of messenger RNAs contain many important cis-regulatory elements that are under functional and evolutionary constraints. We hypothesize that these constraints are similar to grammars and syntaxes in human languages and can be modeled by advanced natural language models such as Transformers, which has been very effective in modeling protein sequence and structures. Here we describe 3UTRBERT, which implements an attention-based language model, i.e., Bidirectional Encoder Representations from Transformers (BERT). 3UTRBERT was pre-trained on aggregated 3’UTR sequences of human mRNAs in a task-agnostic manner; the pre-trained model was then fine-tuned for specific downstream tasks such as predicting RBP binding sites, m6A RNA modification sites, and predicting RNA sub-cellular localizations. Benchmark results showed that 3UTRBERT generally outperformed other contemporary methods in each of these tasks. We also showed that the self-attention mechanism within 3UTRBERT allows direct visualization of the semantic relationship between sequence elements.

## 1. INTRODUCTION

The 3’untranslated regions (3’UTRs) of messenger RNAs (mRNAs) are critical in regulating gene and protein expression at the post-transcriptional level. Cis-regulatory elements located in the 3’UTRs are recognized and bound by trans-acting factors such as RNA-binding proteins (RBPs) and microRNAs, which can modulate mRNA modification, abundance, and subcellular localization [1–3]. A number of computational methods have been developed to predict these cis-regulatory elements, either by matching mRNA sequence with experimentally determined binding motifs, or by taking advantage of evolutionary conservation information, or a combination of both approaches. For example, by comparing closely related genome sequences, Kellis, Xie and colleagues derived putative regulatory elements in yeast and mammals, respectively [4, 5]. More sophisticated methods such as PhastCons and PhyloP incorporate phylogenetic information with hidden Markov models to identify genomic regions that are under evolutionary constraints, which have been widely used in the field [6, 7]. A number of specific computational tools were also developed for specific tasks such as RNA-RBP interaction [8], m6A modification preference [9], and mRNA subcellular localization [10].

In recent years, advances in deep learning (DL) have prompted researchers to develop DL based methods in the study of RNA regulation; these methods include Convolutional Neural Network (CNN) [11], Recurrent Neural Network (RNN) [12], or integration of both architectures [13, 14]. Despite their success, these methods can have limitations such as difficulty to parallelize, or vanishing gradient when processing extra-long sequences. The CNN-based methods are parallelizable, but they often focus on local contextual information and are unable to model long-range semantic dependency [15]. Recent breakthroughs in natural language processing (NLP) models have prompted researchers to explore to use these powerful NLP models in the study of biological sequences. The rationale behind these approaches is that biological sequences such as protein amino acid sequences or regulatory elements follow evolutionary or functional constraints that are similar to the evolution of syntax and gramma of human language, so that NLP models can be applied to these sequences to derive the latent constraints or “grammars” hidden in the biological sequences [16]. Among these modern NLP models, attention-based Transformers and Bidirectional Encoder Representations from Transformers (BERT) were recently adapted to model biological sequences on specific tasks [17, 18]. Ji et al. introduced DNABERT, a Transformer model pre-trained on the whole human reference genome that enabled fine-tuning for specific tasks, such as the annotation of promoters or splice sites [15]. Brandes and colleagues developed a protein-BERT model, which allowed them to predict protein properties such as secondary structure, stability, and remote homology [19]. Elnaggar and colleagues evaluated several auto-regressive models and autoencoder language models, and described an embedding based model called ProtTrans, which established that NL models can be useful in learning protein properties [20]. Similarly, Yamada et al. developed BERT-RBP, which was fine-tuned on eCLIP-seq datasets, allowing fine-tuning on eCLIP-seq data to identify RNA-protein interactions [21]. Recently, NLP models were also successfully applied to predict protein 3-D structures [22, 23].

Motivated by the success of language models on the study of proteins, we herein describe a new method, 3UTRBERT, which uses a language model to identify constrained regions in the 3’UTR of mRNAs. Language models applied on the protein sequences can reveal sequence and structure constraints, while language models on genomic DNAs can help identify potential transcriptional regulatory elements. We hypothesize that language models can be applied to mRNA sequences to help identify regulatory elements important at post-transcriptional level. To the best of our knowledge, there have not been any published studies on deploying Transformer or other NLP models on mRNA sequences. We note that Akiyama and colleagues recently described a method called RNABERT, but it was designed to use embedding to improve non-coding RNA structure alignment instead of functional annotations [24]. Our method, 3UTRBERT, was pre-trained on aggregated 3’UTR sequences of human mRNAs in a task-agnostic manner; the pre-trained model was then fine-tuned for specific downstream tasks such as predicting RBP binding sites (41 RBPs covering multiple CLIP experiments), m6A RNA modification sites, or RNA sub-cellular localizations. For each of these sub-tasks, we benchmarked the performance of 3UTRBERT against state-of-the-art methods. Our results showed that 3UTRBERT generally outperforms other methods for these tasks. We also demonstrated that these identified informative sequence motifs can be visualized by our pipeline as regions of higher attention scores.

## 2. RESULTS

### 2.1 Overview of 3UTRBERT design and applications

**Fig. 1** shows the overall design of 3UTRBERT. The top of the figure shows that 3UTRBERT was first trained with unannotated RNA fragments using self-attention-based Transformer architecture. The correlations among token positions learned in the pre-training stage were aggregated into the final hidden vector of the [CLS] token to provide general and transferable information. This offers the model the flexibility to work with both long RNA sequences and short RNA fragments in the downstream tasks (see **Materials and Methods**). For classification problems involving longer sequences, we freeze the model parameters and extract the contextual embedding inside 3UTRBERT as the effective feature scheme for each base. Meanwhile, 3UTRBERT can also be directly fine-tuned on binary labels of short-region classification tasks in an end-to-end manner. We further demonstrated the flexible configurability and the effectiveness of such knowledge-embedded framework for transcript labelling. 3UTRBERT compared favorably against the latest methods for predicting RBP-RNA interactions, m6A modifications, and mRNA subcellular localization (bottom half of **Fig. 1**). In addition, the interpretable analysis implemented by self-attention mechanism also allowed visualization of the semantic relationship among nucleotides.

**Fig. 1.**
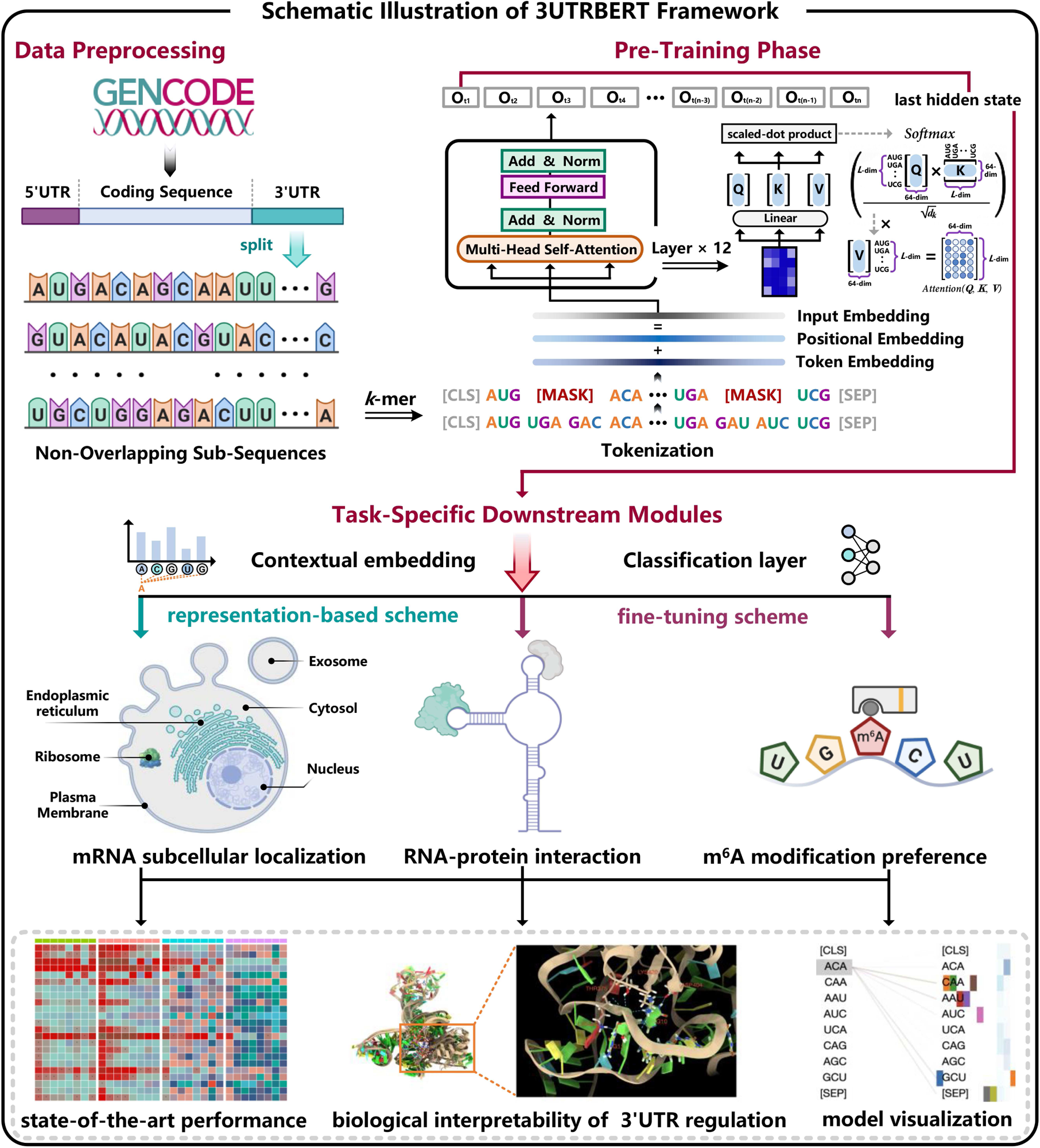
Schematic illustration of 3UTRBERT framework. 3UTRBERT is pre-trained on the 3’UTR sequences of human genes using self-supervised learning. Unlabeled mRNA sequences are segmented by sequential 3-mer tokens and subsequently fed into a Transformer architecture to learn general-purpose representations using a masked language model (LM), i.e., reconstructing the masked tokens from other tokens thus learning the syntax and grammar hidden in the 3’UTR. The pre-trained model is then fine-tuned on several downstream prediction tasks, including predicting RBP-RNA interactions, m6A modification sites and subcellular localization.

### 2.2 3UTRBERT improves detection of RBP binding sites

RNA-binding proteins (RBPs) typically recognize specific sequence motifs facilitated by local RNA structure features on the mRNA transcripts [25]. We hypothesize that the sequence windows with high attention scores were enriched with these RBP binding sites. As described in **Methods**, we downloaded and processed protein-RNA crosslinking sites for 41 RBPs and used these experimentally determined binding sites to fine-tune the Transformer model to predict RBP binding sites. We next benchmarked the performance of 3UTRBERT with several other state-of-the-art computational methods: a CNN and RNN-based model, a graph neural network (GNN)-based model, and a Transformer-based model (**Fig. 2a** and **2b**). iDeepE combines local multi-channel CNNs and a global CNN to predict RBP binding sites from sequences alone [26]; DeepCLIP uses one-hot RNA encoding to represent RBP binding sites and constructs a shallow neural network composed of CNN and LSTM layers [13]; RPI-Net employs a GNN framework and adds base-pairing information to enhance prediction accuracy [27]; Graphprot2 employs graph convolutional neural network (GCN) with RNA secondary structure information to infer protein-RNA binding preferences [28]. Despite promising results, these methods typically only work on localized mRNA regions of 100 nucleotides (nts) long, and the deep neural network architectures often lack interpretability and require longer training time. Besides these deep learning-based approaches, RNABERT uses a Transformer module to generate informative embedding for each nucleotide from non-coding RNAs; however, this method was intended for predicting RNA structures instead of detecting regulatory elements [24]. Another method, BERT-RBP, only utilizes fine-tuning to infer RNA-protein interactions on eCLIP-seq data and does not use RNA sequence properties in pre-training stage [21]. For our benchmark studies, we included iDeepE, DeepCLIP, RPI-Net, GraphProt2, RNABERT and BERT-RBP.

**Fig. 2.**
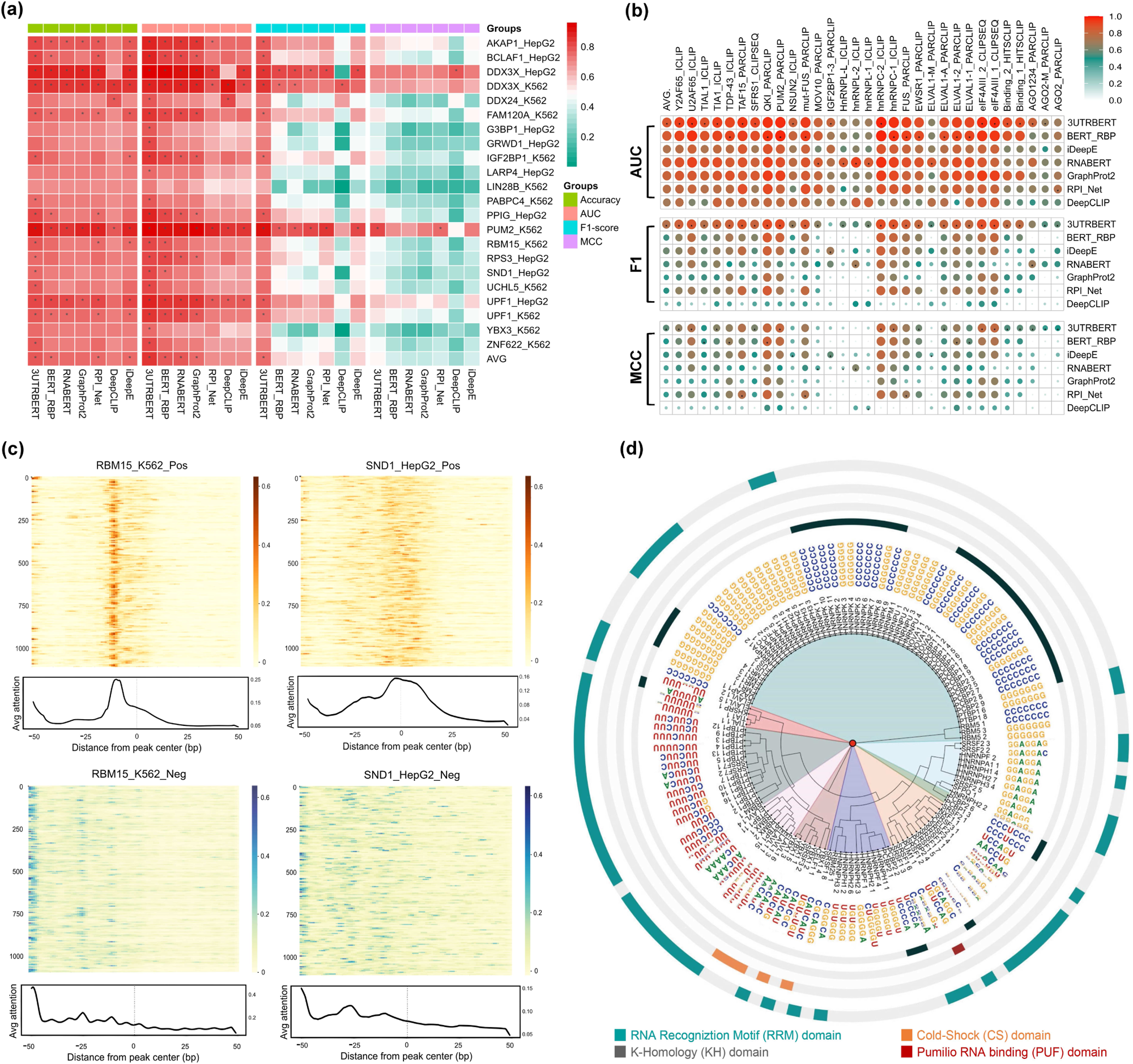
**(a)** Performance comparisons between 3UTRBERT and other methods on predicting RBP-RNA interactions (22 RBPs); prediction accuracy is color coded. **(b)** Performance comparisons between 3UTRBERT and other methods on 19 additional RBP CLIP datasets; prediction accuracy is coded by size and color of the bubbles. (**c**) Landscape of attention scores surrounding RBP-RNA binding sites and the negative control sites; RBM15 in K562 cell line and SND1 in HepG2 cell line are shown as examples. (**d**) Radial tree plot showing the congruent clustering of RBP-binding motifs and RNA-binding domains.

We used 5-fold cross-validation to test the performance of the predictive methods (please see **Methods** for details). The following metrics were used to quantify the performance of predictions: (i) AUROC (Area Under the Receiver Operating Characteristic), (ii) F1-score (the harmonic mean between precision and recall), (iii) MCC (Matthews Correlation Coefficient), and (iv) ACC (accuracy). The formal definition of these metrics can be found in **Supplementary Note 1**. We first conducted benchmark studies on 22 RBPs that were described in an early study [29], and reproduced the results of other predictors as described in the original publications using default parameters. **Fig. 2a** shows that 3UTRBERT generally outperformed other methods (AUROC = 0.836, F1 = 0.751, MCC = 0.503, and ACC = 0.785), while the second best results were AUROC=0.788 (by BERT_RBP), F1=0.565 (by iDeepE), MCC = 0.413 (by iDeepE) and ACC=0.758 (by iDeepE) (**Supplementary Table 1**). We next compared the performance of these methods on 19 other RBPs described from a more recent publication [30], and ensured that these 19 RBPs have no significant sequence homology with the 21 RBPs we used for fine-tuning. **Fig. 2b** shows 3UTRBERT also outperformed other methods on these RBPs (more details are in **Supplementary Table 2**). For example, for two proteins, Ago and IGF2BP, 3UTRBERT achieved the best mean performance with AUROC of 0.791± 0.008 and 0.805± 0.003, respectively. We note that, although BERT-RBP was specifically designed to study RBP-RNA binding, it didn’t achieve the best performance, perhaps due to lack of pre-training steps [21]. RNABERT also did not perform well since it was designed for the purpose of predicting RNA structures [24] (**Supplementary Fig. 1**). Other deep-learning or graph-based approaches also did not perform as well due to likely architectural complexity and insufficient representational capabilities. Overall, these results demonstrated that the representation learning approach implemented in 3UTRBERT was an effective approach in finding RBP binding sites located in the 3’UTRs.

**Fig. 2c** shows the landscape of attention scores for RBM15 and SND1. For positive samples, the nucleotides adjacent to the cross-linking sites tend to have the highest attentions scores in the 100-nucleotide window. However, the attention scores of negative samples did not have such enrichment. We next investigated whether these continuous high attention regions are enriched with previously known RBP binding motifs. Specifically, we used different cutoffs to select regions of high attention scores and extracted enriched sequence motifs. These sequence motifs were subsequently converted to positional weight matrices (PWM) and compared with known RBP binding motifs collected in the ATtRACT database [31]. By using the TOMTOM search tool [32] with an FDR q-value <0.01, 133 documented motifs were discovered and matched to the candidate motifs discovered by attention scores. We further systematically visualized the hierarchical clustering of all validated motifs (**Fig. 2d**). We followed the same scheme as a previous publication [33] and showed that RBPs involved in similar RNA regulatory pathways were generally grouped together within the canonical RNA binding domains (RBD) [34]. For example, HNRNPK clearly prefers to cytosine (C)-rich motifs within 3’UTR of K-Homology (KH) domain to stabilize target mRNAs, the dysregulation of which is implicated in cancer progression [35].

We next carried out RNA-protein docking and molecular simulation experiments to test the physical interactions between RBPs and the RNA motifs identified by 3UTRBERT. Due to computational complexity, we selected RBM15 and SND1 for this study since these are well-known RBPs implicated in critical cellular processes such as m6A modifications and tumor angiogenesis [36, 37]. We used the experimentally determined crystal structure for RBM15 (PDB ID Q96T37) and the predicted protein structure for SND1 (AlphaFoldDB, 5M9O) [38]. For each of these RBPs, we took the sequence motif of the highest attention score, GGCCCG for RBM15 and UUCCAGG for SND1. We investigated several 3D RNA structure prediction tools and used DeepFoldRNA [39]; we confirmed that the structure predictions converged and returned converged final structures. We next used the ClusPro web server to perform RNA-protein docking to generate high-resolution 3-D RBP-RNA complexes [40]. ClusPro uses PIPER to generate docking samples, which implements a Fast Fourier Transform (FFT) algorithm with structure-based pair-wise interaction terms. Specifically, it uses shape complementarity, electrostatic, and de-solvation terms to produce the near-native binding complexes. ClusPro further clustered the 1,000 most energetically favorable samples by root-mean-square deviation (RMSD), and then performed structure refinement on the cluster centers using Charmm energy minimization, ultimately returning top 10 binding complexes [41]. After acquiring these samples, we filter for those involving interactions with the RNA motifs identified by our method. For RBM15-RNA complexes, the median PIPER energy score for top 10 docked complex was -1792.6, -1839.4 for top five complexes; for SND1-RNA complexes, the median PIPER energy score for top 10 docked complex was -1269.1, -1310.1 for top five complexes. We note that these numbers were calculated from scoring functions, which were proportional to free energies but did not have actual units.

After docking experiments, we further conducted molecular dynamic (MD) simulation on docked RBP-RNA complexes after filtering to further refine the RBP - RNA complexes and evaluate interactions between RBP residues and RNA nucleotides. **Fig. 3a** and **3b** show the average structures for the final 50 nanoseconds of simulation (more details in **Supplementary Note 2**). We observed formation of strong phosphate contacts, accompanied by supplementary base contacts, further supporting the reliability of the predicted RNA-protein interactions. In the case of RBM15, the predicted RNA motif was located at a loop forming a pseudoknot; the side chain residues of the first three beta sheets of RBM15 flipped the nucleotides on the motif outwards, forming two notable hydrogen interactions: (ASP404 - G10) and (LYS450 - G11), and the phosphate interactions were also observed for (THR375, G11) and (LYS420, G10, G11). For SND1, the bound RNA motif was situated on a hairpin loop; the side chains on the second to fourth beta sheets of SND1 were inserted into the hairpin, forming multiple hydrogen bond interactions with the bases, specifically (GLU744 - A58), (ASP765-G60), and (TYR766 - G59). Interestingly, the predicted electrostatic interactions with nucleic acids GGA (58-60) have been previously reported as a binding site of SND1 for m6A modification site recognition [42]. Overall, our docking and MD experiments, despite with only on two RBPs, showed that the RNA motifs discovered by 3UTRBERT were biologically meaningful and could generate insights on interactions between RBP and target RNA sequences.

**Fig. 3.**
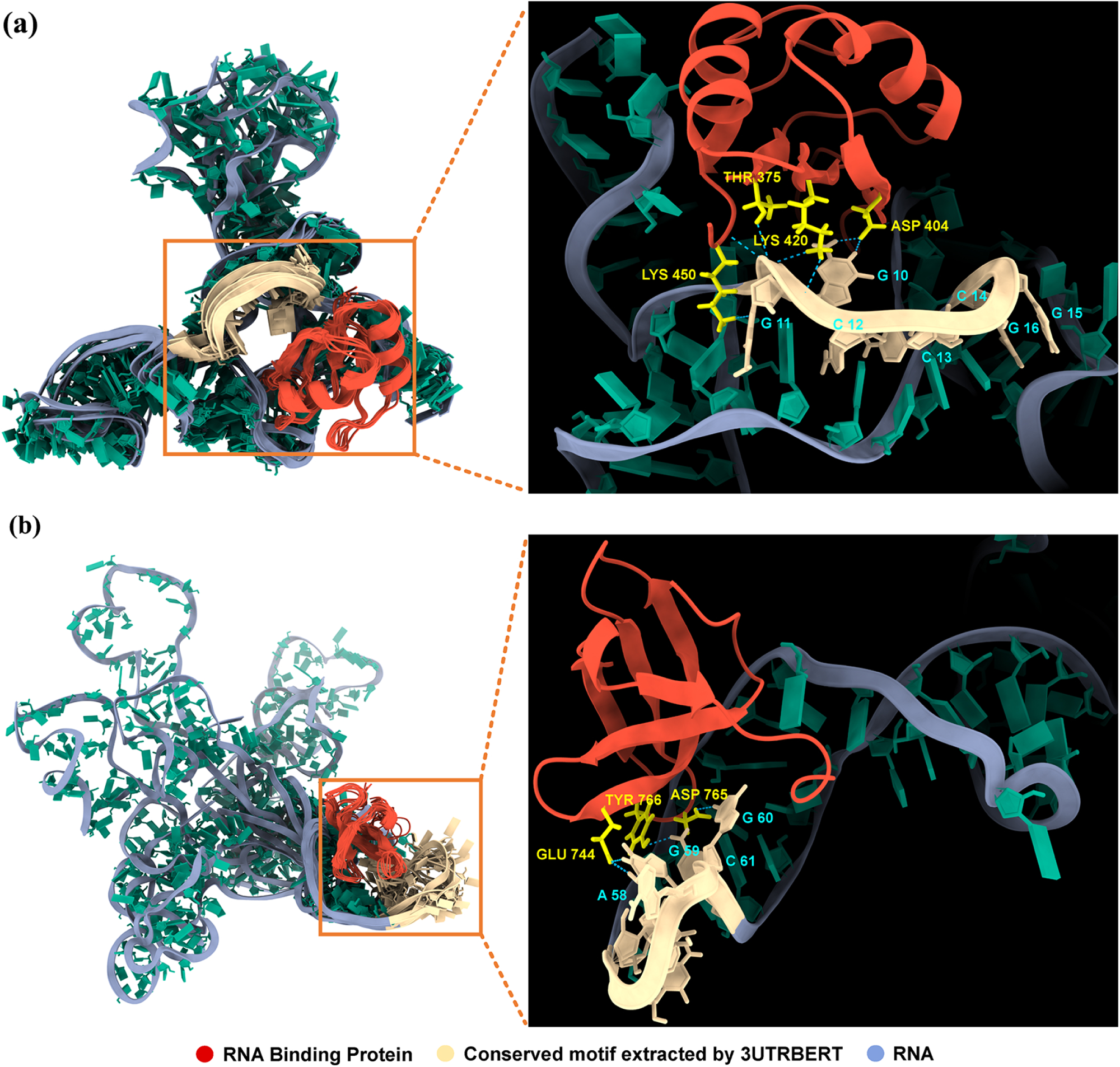
(**a**) Predicted complex structure between RBM15 and a target RNA sequence (GGCCCG), after protein-RNA docking and all-atom molecular dynamics simulations. Hydrogen bonds between side chain residues of RBM15 and RNA are observed: Thr375-G11, Asp404-G10, Lys420-G10/G11, and Lys450-G11. (**b**) Predicted complex structure between SND1 and a target RNA sequence (UUCCAGG). Multiple hydrogen bonds are observed: Glu744-A58, Asp765-G60, and Tyr766-G59.

### 2.3 3UTRBERT improves identification of m6a modification sites

Inside cells, RNA transcripts undergo extensive covalent modifications. Among the estimated dozens of types of modifications, m6A (N6 methyl adenosine modification) is the most prevalent and most studied. It is reported that over 70% of the mammalian mRNA transcripts inside cells undergo m6A modifications and m6A is implicated in many important developmental processes and human diseases [43–45]. Many of these modification sites are evolutionarily conserved, although there are debates on the strength of these evolutionary constrains [46]. We hypothesize that evolutionary conservation and sequence contexts of these m6A sites would allow 3UTRBERT pick them out from background sequences. Towards this goal, we fine-tuned the 3UTRBERT model to predict potential m6A modification sites. As described in Methods, the ground truth m6A sites were downloaded from m6A-Atlas database [47].

We benchmarked 3UTRBERT with several other m6A sites predictors, including machine learning (ML) based methods (SRAMP)[48], WHISTLE [49], and iMRM [8]) and a deep learning (DL) based method (DeepM6ASeq [50]). The results are illustrated in **Fig. 4a** and **Supplementary Table 3**, showing that 3UTRBERT mostly outperformed other methods in four different metrics and in 9 different cell lines. SRAMP and WHISTLE used hand-crafted features and generally didn’t perform as well as other methods, whereas iMRM depended heavily on cross-species conservation. DeepM6ASeq was based on a hybrid deep learning network architecture and generally achieved the second-best performance, just below 3UTRBERT. **Fig. 4b** showed two examples from two cell lines (A549 and HCT116) as depicted by visualization tool kpLogo [51], showing high attention scores associated with experimentally determined m6A site from m6A-Atlas database. Additional examples can be found in **Supplementary Fig. 2**. Interestingly, we found that our learned consensus motifs (highlighted with red-colored boxes) also closely matched the consensus m6A motif RRACH (blue-color window) (where A = m6A, R = purine, and H = A, C, or U) [43]. The Transformer architecture in 3UTRBERT allowed us to visualize the transition and convergence of attention scores in a sequence. As an example, **Fig. 4c** provides a bird-eye view across 12 attention heads on an RNA sequence. The sequence was represented as a series of 41 3-mer tokens, in addition to [CLS] and [SEP]. 3UTRBERT correctly identified two important regions (red boxes with self-attention converged), which were known m6A sites. By adjusting thresholds, we were able to uncover the hidden semantic relationship embedded in the crucial token “ACU”, which aggregated the contribution attentions from various short regions.

**Fig. 4.**
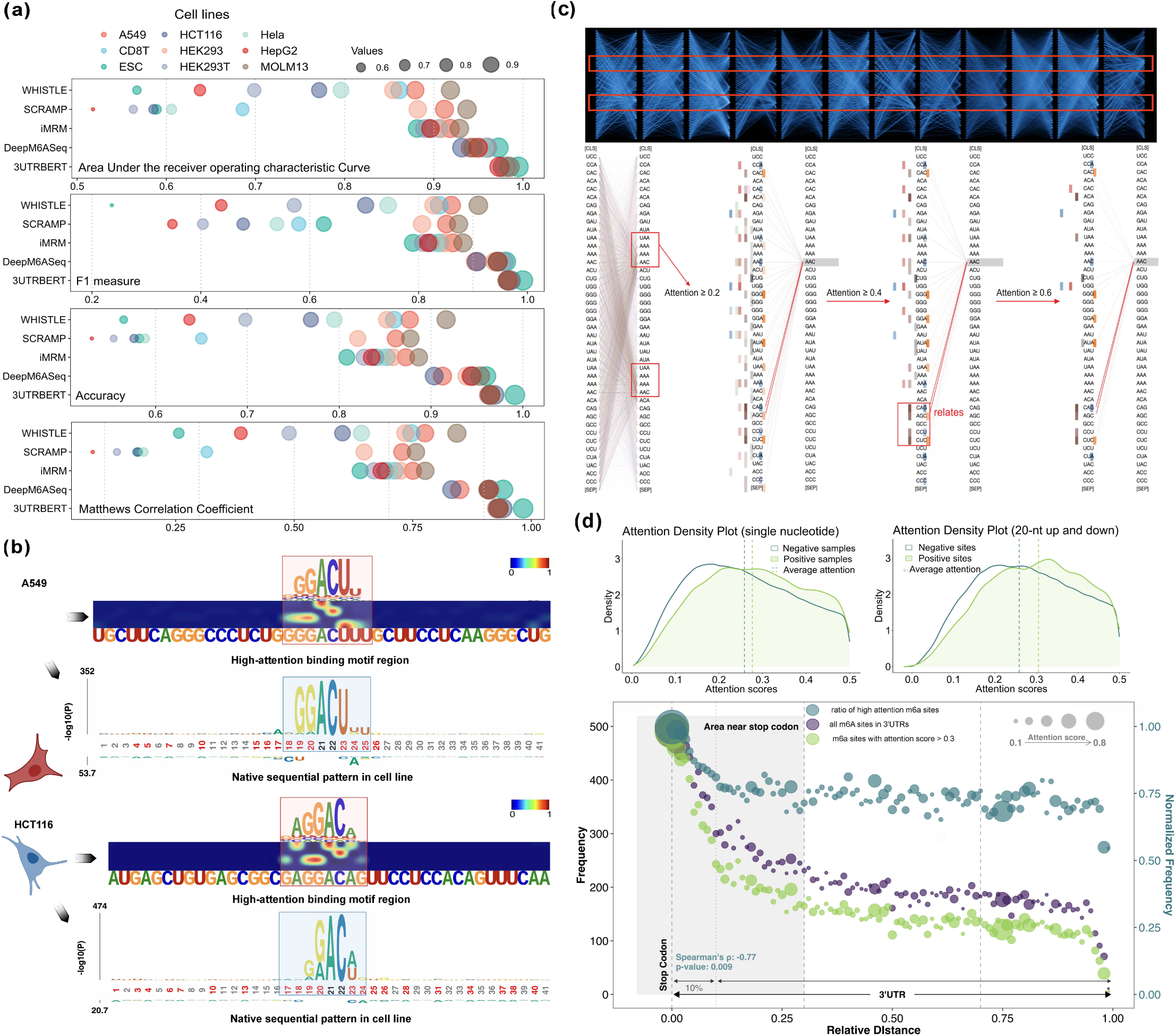
(**a**) Bubble plots comparing prediction performance between 3UTRBERT and other methods. Each panel corresponds to a statistical metric: AUROC, F1 measure, Accuracy, and Matthew’s correlation coefficients. (**b**) Two examples of predicted m6A sites with high attentions scores and consistent with well-defined RRACH m6A motif. (**c**) A bird’s-eye view of global attention scores throughout entire architecture (top), showing 3UTRBERT correctly self-focused on two converged regions (red boxed) corresponding to known m6A modification sites. The details of contextual relationship within *k*-mer level are shown in Attention-head plot (bottom). (**d**) (top: distribution of attention scores of experimentally determined m6A sites (positive) and negative control sites, at single nucleotide level and within 40 nucleotides window. (bottom) Distribution plots showing the predicted m6A sites are enriched near stop codons (see text).

We next investigated whether globally the known m6A modification sites had higher attention scores than the background. **Fig. 4d** (top) plots the distribution of the attention scores of the experimentally determined m6A sites and negative control positions (please refer to **Materials** for the choice of negative set), which shows that the m6A sites have overall higher attention scores (p-value < 1.19e-55, Mann Whitney U test). We investigated how the attention scores were spatially distributed among 3’UTRs, by scaling 3’UTR sequences into the same length and calculated the frequency of high attention scores along the sequence; for example, relative distance of an m6A site at position 10 on a 50 nucleotides-long 3’UTR was calibrated as 0.2 (10/50). **Fig. 4d** (bottom) shows that m6A sites were enriched in the region closer to stop codons. The purple bubble curve in **Fig. 4d** (bottom) shows that the number of m6A sites decreased with growing distance from the stop codon. We next asked whether the m6A sites closer to stop codons also have higher attention scores, i.e., being stronger modified sites. We applied a threshold of 0.3, which is the average attention score over aggregated 3’UTRs, and retained only those m6A sites with attention scores higher than 0.3 (green circles). It is clear that higher attention m6A sites are also accumulated near the stop codons, which has been previously shown [47, 52]. We next calculated the fraction of high-attention m6A sites over total number of m6A sites in sliding windows (dark blue circles), which shows that indeed the high attention scores tend to be enriched closer to stop codons, i.e., negatively correlated with the distance to stop codons (Pearson correlation =-77% and p-value < 0.009). These observations demonstrated that indeed the attention based 3UTRBERT can effectively predict and recover experimentally determined m6A sites [43, 44].

### 2.4 3UTRBERT helps to predict mRNA subcellular localization

The 3’UTR of mRNA transcripts contain sequence elements that can help direct transportation of mRNA transcripts to specific subcellular compartments [53]. These experimentally determined localization data have been catalogued in databases such as RNALocate and RNALocate v2.0 [54, 55](see **Materials and Methods**). Several computational algorithms have been published, aiming to predict mRNA localization from RNA sequence or structure signatures. These methods include DM3Loc [10], RNATracker [56], mRNALoc [57], and iLoc-RNA [58]. Briefly, iLoc-mRNA and mRNALoc use Support Vector Machine (SVM) with hand-craft features. RNA-Tracker integrates sequence and secondary structure information using a neural network framework. DM3Loc uses multi-head self-attention mechanism in predictions; notably it also predicts multi-label subcellular localizations, i.e., allowing an mRNA transcript present in multiple subcellular compartments [10]. Despite promising results, these methods still have limitations, as it is difficult to model long-range dependencies among nucleotide positions by the widely used sparse positional encoding scheme. The major innovation of 3UTRBERT versus the other methods is that it creates an informative coding scheme by generating context-sensitive embedding with self-learning. Adopting a framework similar to DM3Loc, we trained 3UTRBERT for the sub-task of predicting RNA localization and benchmarked against four other methods. **Fig. 5a** summarizes the performance metrics of AUROC (Area Under the Receiver Operating Characteristic) and AUPRC (Area Under the Precision-Recall Curve). Only the compartments predicted by all five methods are listed; detailed results can be found in **Supplementary Table 4**. The left panel on **Fig. 5a** showed that 3UTRBERT had the best performance on both AUROC and AUPRC, with the DM3Loc coming in second in performances.

**Fig. 5.**
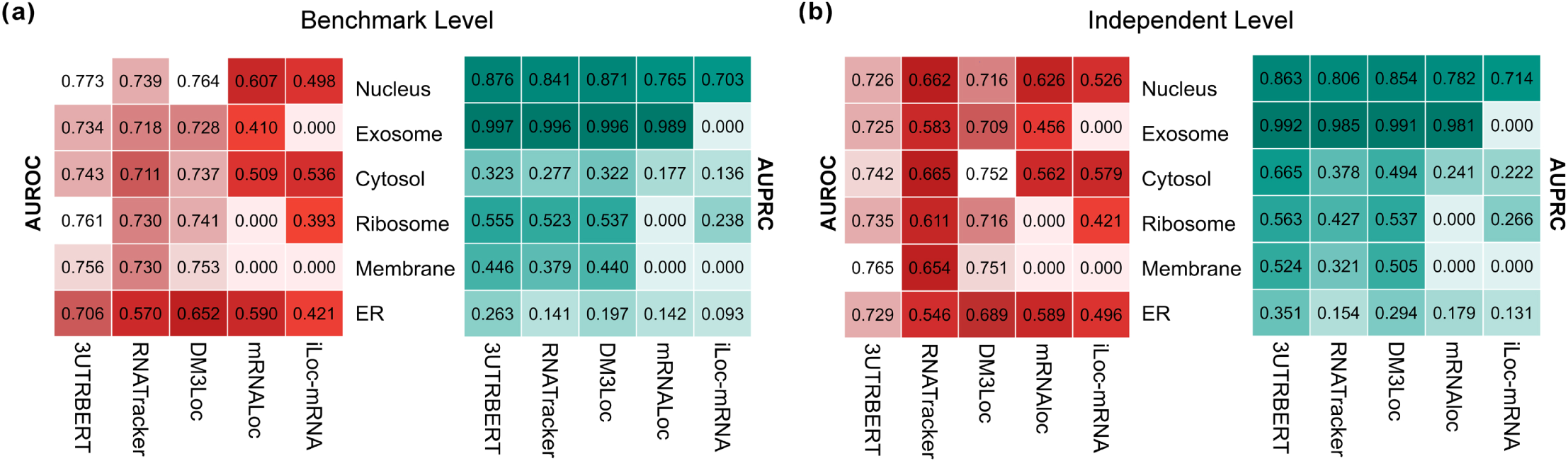
(**a**) Comparison of prediction performance between 3UTRBERT and other methods on mRNA subcellular localizations. Localization data from DM3Loc was used. (b) Prediction performance on a newer independent set of localization data from RNALocate 2.0.

To further evaluate the robustness of 3UTRBERT in predicting mRNA subcellular localizations, we conducted another benchmarking experiment on an independent dataset from RNALocate v2.0 database [55], containing a collection of mRNA sequences that were excluded from all previous datasets. Employing the models described above, we directly fed the test data to the model to make predictions (**Fig. 5b** and **Supplementary Table 5**). 3UTRBERT also achieved the best performance, achieving AUROC of 0.737, and mean average area under Precision-Recall (AUPRC) of 0.66, over six compartments. The second-best results were AUROC = 0.722 and AUPRC = 0.613 (by DM3Loc). We then calculated the average prediction metrices using three different coding schemes (sequence with embedding, sequence with attention score, and sequence only) of six subcellular localizations for comparison (**Supplementary Fig. 3**). The embedding and attention based approaches achieved better performance (73.74% ± 0.009, 73% ± 0.014) than sequential information alone (72.22% ± 0.014) on AUROC value, suggesting that the contextual feature from deep representation learning helped to enhance the original model.

### 2.5 3UTRBERT can derive semantic contextual information from biological sequences

The 3’UTRs of eukaryotic mRNA transcripts contain important sequence motifs that are under evolutionary and functional constraints, which may have evolved in a manner similar to the evolution of syntax and grammars of human languages. This allows us to use advanced nature language models, especially attention mechanisms, to infer and visualize the contextual information between these motifs. Such contextual information can be extracted by self-attention mechanism within Transformer model to weigh and capture relationships between different tokens in the mRNA sequences. The self-attention mechanism operates by assigning attention scores to each token based on its dependencies on other tokens, thus helping 3UTRBERT capture the intricate patterns between different elements within the 3’ UTRs, even if they are separated by several tokens. To search for the most optimal *k*-mer size to balance the computational time and the representation capability, we first tested four types of *k*-mer (k = 3, 4, 5, 6) with 1-stride settings. **Fig. 6a** displays the evaluation results before and after training at different *k*-mer size for the 22 RBP eCLIP results (see **Section 2.2**). For example, the top left panel compares the prediction performance, as measured as AUROC, after the pre-training stage (shown on the horizontal axis) and after the fine-tuning stage (shown on the vertical axis). It is clear that, for all the *k*-mer size tested, the task-specific fine-tuning always improved the prediction performance compared to the task-agnostic pre-training stage. We observed that *k*-mer=3 had the best prediction performance with averaged AUROC = 0.8178 after pre-training and AUROC = 0.8364 after fine-tuning. We also explored fine-tuning directly on the scratch model, i.e., skipping the pre-training phase and directly training the model on a particular task with randomly initialized parameters. As expected, the results and stability of all *k*-mers (0.7208 ± 0.005, 0.6866 ± 0.007, 0.6797 ± 0.009, and 0.6662 ± 0.012) dropped significantly (**Supplementary Fig. 5**). This shows that the pre-training stage learned from a large corpus of unannotated RNA sequences and captured valuable contextual knowledge that can be leveraged for various downstream tasks.

**Fig. 6.**
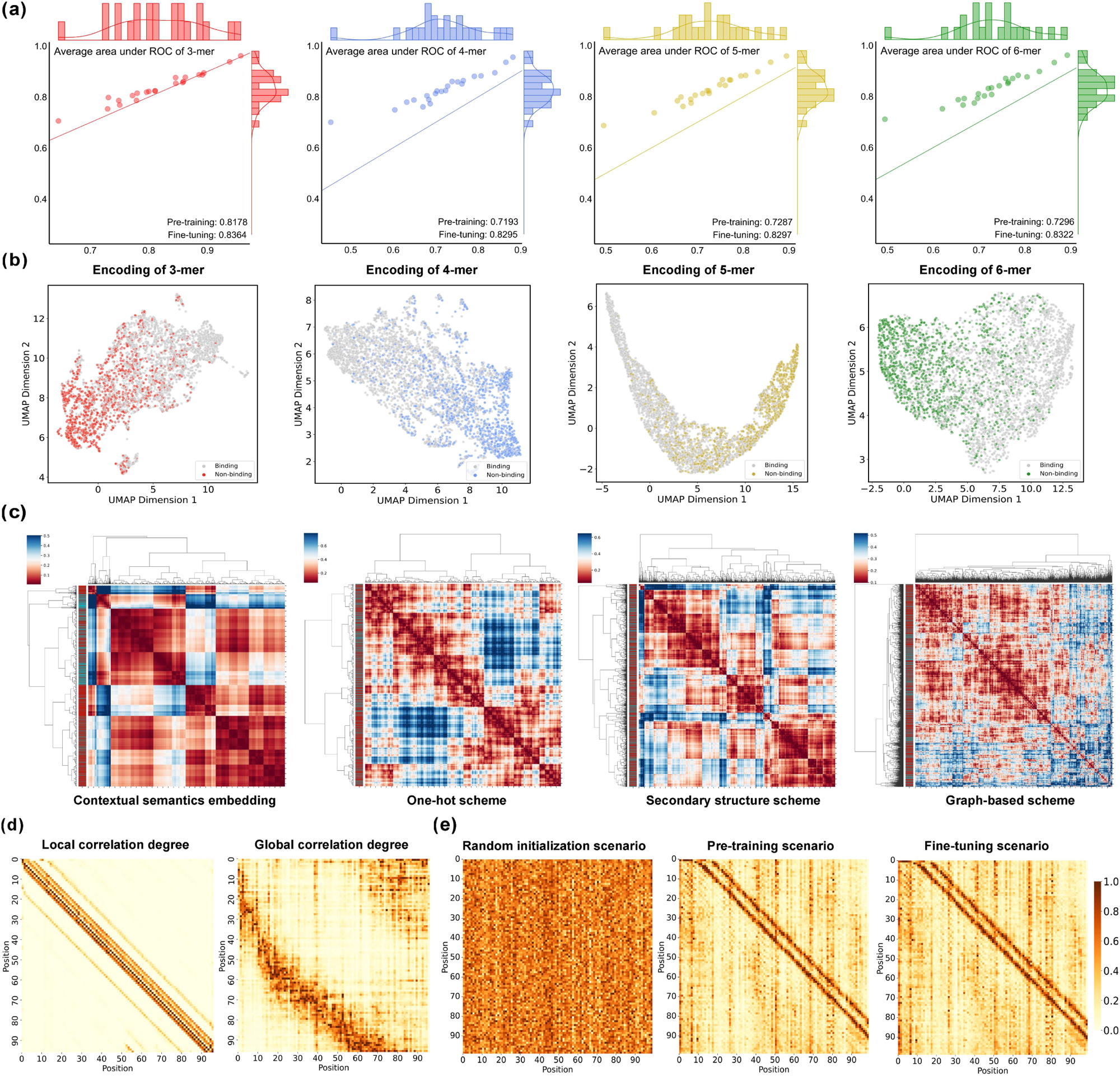
(**a**) Scatter plots of AUROC scores after pre-training (X-axis) and after fine-tuning (Y-axis) for 22 RBP eCLIP datasets. Different *k*-mer sizes are shown separately. Fine-tuning has generally improved the prediction performance. (**b**) UMAP projection of the embedding vectors generated by 3UTRBERT for RBP binding sites (positive) and negative controls; different *k*-mer sizes are shown. (**c**) Clustering of positive RNA sequences bound by RBM15 as encoded by four different coding schemes: 3UTRBERT, one-hot scheme, secondary structure, and a graph-based embedding. (**d**) Attention maps illustrating that 3UTRBERT can extract both local and global contextual semantics. (**e**) Illustration of how information is extracted at scratch, pre-training and fine-tuning stage.

We next tried to project and visualize the feature space of the positive and negative (bound and unbound) RNA sequences as decoded by 3UTRBERT. Taking RBM15 as an example, **Fig. 6b** uses UMAP (Uniform Manifold Approximation and Projection) to visualize the input RNA sequences as two-dimensional projections [59], using different *k*-mer sizes. Each data point represents a sample input RNA sequence (100 nucleotides long), predicted positive (bound) sequences are painted in color while negative (unbound) sequences are in grey. These figures clearly show that the embedding generated by 3UTRBERT very effectively separated the positive and negative samples.

We next investigated whether the language model performs better in clustering input RNA sequences than other encoding schemes. **Fig. 6c** compares the 3UTRBERT token embedding scheme with one-hot scheme, secondary structure, and a graph-based embedding scheme on RNA sequences bound by RBM15 as input. In one-hot scheme, each nucleotide is represented as a binary vector according to base type; in secondary structure scheme, each nucleotide is represented as one of the five structure categories (E for external loop, H for hairpin loop, I for internal loop, M for multi-loop, and P for paired); in graph-based embedding, each RNA sequence is represented as a graph where nucleotides are nodes and their interactions or spatial proximities are edges. We can see that the contextual semantic embedding by 3UTRBERT gave stronger and cleaner clustering of positive bound sequences.

One advantage of attention-based nature language models such as 3UTRBERT is that they can capture both local and long-range sequence semantics or interactions. Such relevance can be visualized with attention map, a high-resolution perspective on RNA sequence analysis. The degree of correlations along the sequences illustrates attention strength between respective positions. By contextualizing RNA sequences both locally and globally (as shown in **Fig. 6d**), 3UTRBERT can help extract correlations or interactions between different elements within 3’ UTRs, which are important to understand regulation at post-transcriptional level. We next portrayed the complete evolutionary processes of hidden semantic relationships (**Fig. 6e**). As we can see, the information is cluttered during the scratch phase, yet by sufficient iterations of the training process, the complex and hierarchical properties are uncovered, and the following fine-tuning operation further enriches the contextual information of each nucleotide on the task-specific data.

Unlike static word embedding techniques that generate fixed vector values for the same token, the Transformer model derives different vector representations based on dynamic changes in its surrounding context. However, deep learning models, including BERT, are often viewed as “black boxes” since it can be challenging to understand why they make certain predictions. Using a novel visualization system Dodrio [60], we linked attention weights with semantic contextual knowledge in order to provide a clear understanding of the learning process of 3UTBERT. We randomly selected two homologous RNA sequences bound by RBM15 and created **Fig. 7a** to visualize the dynamically generated semantic information. Specifically, these two sequences were selected by BioPython [61] based on a high global alignment score, and then aligned locally using BLAST [62]. In **Fig. 7a**, the “Attention Head Overview” displays the relative importance of all attention heads for the input RNA sequences. Each head was comprised of weights incurred from 3-mer tokens when calculating the next representation of the current token, and then formed a Radial Layout attention map. For the identical position of the attention head, the final hidden vector of the [CLS] token aggregated different semantic relationships, resulting in distinct decisions for downstream classification tasks. More importantly, the distribution of attention heads with high contribution scores increased by the number of layers and peaked at the last layer, which is consistent with the UMAP classification in each layer of the 3UTRBERT model (**Supplementary Fig. 6**). In the context of RNA sequences, lower layers are closely associated with local features such as specific nucleotide patterns or motifs, while higher layers gradually integrate the contextual information related to identifying functional sites. We next used the SHAP tool to visualize the impact of various regions on the RNA sequence (*k*-mers) on model predictions [63]. The regions with high SHAP values (red color), as visualized in **Fig. 7b**, are correlated with functionally important regions of the RNA, i.e., RBP binding motifs. The conserved GC-rich semantic regions have the same positive effect on model predictions for different RNA sequences. This strongly demonstrated that the knowledge-embedded framework of 3UTRBERT can learn and understand functional semantics of the biological sequences and can be easily extended to other sequence labelling tasks.

**Fig. 7.**
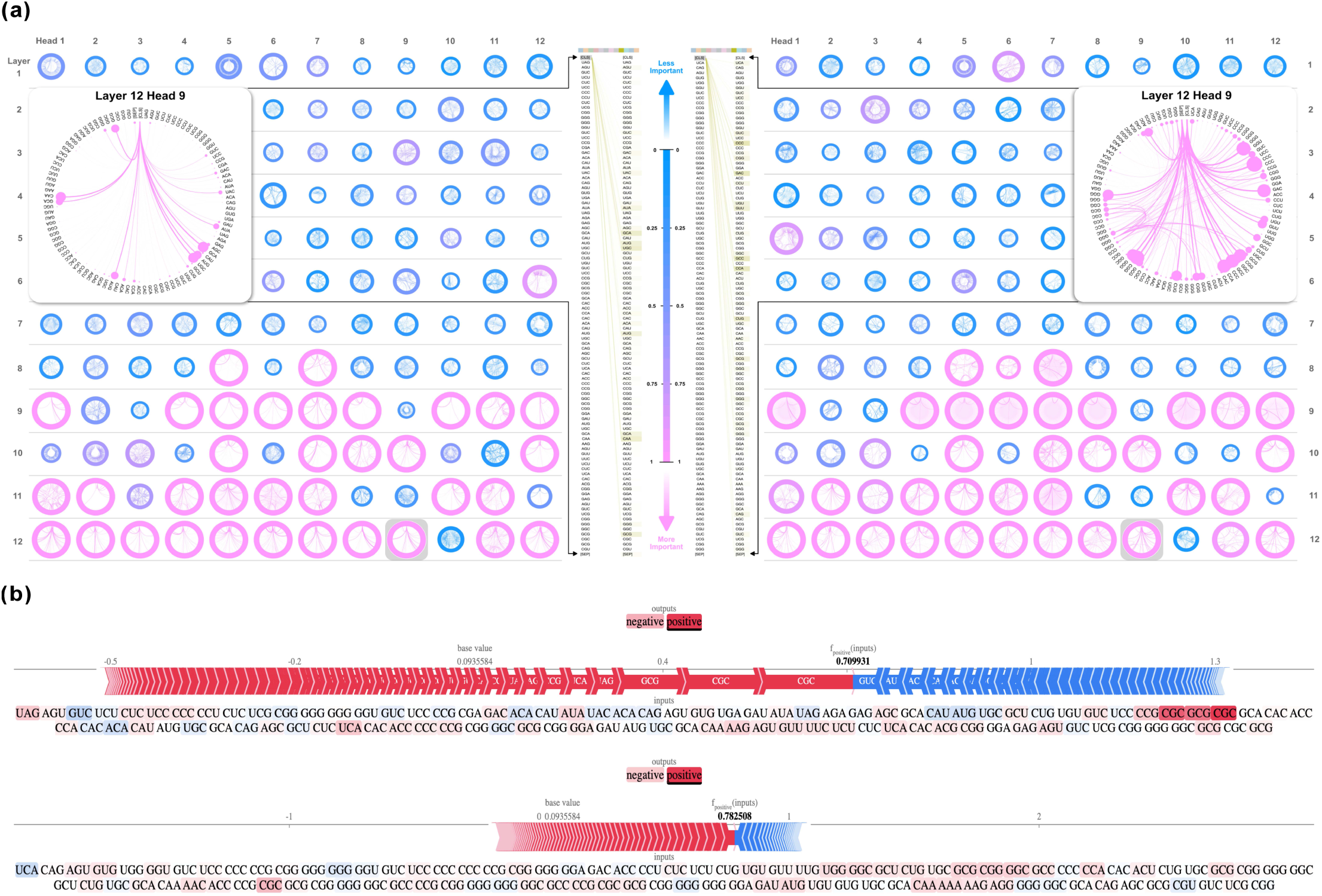
(**a**) Attention Head Overview provides the contribution score in each attention head for the input sequences. The dynamic representation-based 3UTRBERT shares different contextual information for homologous sequences. (**b**) Textplot visualizes the importance of individual tokens in 3UTRBERT. A token with a strong positive SHAP value is highlighted in red color, indicating a positive prediction from the language model, while a negative SHAP value is marked with a blue color, indicating a decrease in the model’s prediction.

## 3. DISCUSSION

The 3’ untranslated regions (3’UTRs) of mRNAs are essential for post-transcriptional gene regulation, which are conferred by evolutionarily conserved (and in some cases less conserved) sequence or structure elements. We hypothesize that these sequence motifs have evolved in a way like human languages, following specific grammar and syntax to confer required biological roles [16]. Natural language models have recently been applied to predicting protein 3-D structures with great success [22, 23]. Motived by this, we developed 3UTRBERT, which is a Transformer-based framework that can be pre-trained on large body of 3UTR sequences, learn the semantics and grammar, and be further fine-tuned for specific biological tasks such as predicting protein-RNA binding, RNA modifications, and RNA sub-cellular localizations. Benchmark studies against other state-of-the-art methods for these individual tasks demonstrated that 3UTRBERT generally outperformed or performed equally well.

We believe the 3UTRBERT model, and other similar nature language-based models, have the following major advantages. Once trained on a large amount of data and learned the intrinsic structure and syntax of the data, it is quite flexible to adapt to specific subtasks with minimum fine-tuning. The pre-training step can be computationally expensive, but it only needs to be performed once. The pre-training step is also un-biased without the need of labeling, which is in contrast to most other probabilistic or rule-based approaches (the fine-tuning step does require labeled datasets).

There are several avenues in which 3UTRBERT can be improved or extended. We only trained the model on 3UTRs from human genes; it would be intriguing to include 3UTRs from other evolutionarily related mammalian or vertebrate species. It is reasonable to assume that increased training data will improve prediction accuracy and reveal more subtle sequence features. We will need to experiment with different threshold to investigate the necessity and effect of including or removing 3’UTR from orthologous genes from related species. It is also worthwhile to investigate whether 3UTRBERT or natural language models can effectively discover lineage specific sequence signals, which could be “buried” when large corpus of sequence data from many species is used in the pre-training stage. Lastly, there is a still relatively paucity of experimentally determined functional data on RNAs, except for in vitro RBP-RNA binding affinities and selected in vivo RBP-RNA binding events. With RNA related data becoming increasingly available in the coming years, we expect 3UTRBERT or other similar nature language-based models will become more effective in training and discovering informative RNA sequence features.

## 4. MATERIALS AND METHODS

### 4.1 3’UTR functional data benchmark datasets

We collected annotated functional elements in the 3’UTR of human mRNA transcripts and used these elements as ground truth to evaluate and benchmark the performance of 3UTRBERT. Some of these elements were extracted from previous published studies. All the datasets in this study can be downloaded from https://figshare.com/articles/dataset/3UTRBERT_dataset_availability/22845644.

### 4.2 Data preparation for pre-training

For task-agnostic pre-training phase of 3UTRBERT, we downloaded the human mRNA transcript sequences from the GENCODE website (GRCh38.p13, Release 40) [64], which consists of 108,573 unique mRNA transcripts. For genes with multiple transcripts, we chose the longest primary transcript. On average, each of these transcripts contain 3,754 nucleotides (median 3,048 nts), 1,657 nts for coding region (CDS, median 1,215 nts), 276 nts for 5’UTR (median 175 nts), and 1,227 nts for 3’UTR (median 631 nts). BioPython was used in sequence manipulation and alignment [61]. In our model, we used sequence window length of 510 nts since current BERT model has a maximum sentence length of 512 characters (a special token is added at each end). Each 3’UTR sequence was split into non-overlapping windows of 510 nts long, the remaining sequence was back-padded to the same length. As we describe below, we limited the study to 3’UTR of mRNA sequences to avoid codon constrains in the CDS region, and to reduce increased complexity of the entire mRNA transcripts.

### 4.3 Collection of RBP binding sites

We collected data from 31 published and curated CLIP (crosslinking and immunoprecipitation) experiments corresponding to 19 RBPs (RNA Binding Proteins), which provided nucleotide resolution binding sites of RBPs [26, 30, 65]. These CLIP datasets were processed using the same pipeline, and sequence windows of 101 nucleotides were used in peak calling. Positive datasets (binding sites) were derived from nucleotide positions with the highest read-counts; binding sites were required to be at least 15-nucleotides apart to minimize redundancy and prevent neighboring sites. The negative control sets were randomly sampled from the genes without detectable interactions in any of the 31 experiments. In each experiment, a total of 4,000 crosslinked sites were randomly drawn for training purpose, and 1000 samples each were drawn for optimization and model validation. To assure separation between training and testing processes, the independent testing set containing 1000 sequences were sampled only from genes unused in model training [30]. In addition to these 19 RBPs, we also collected a dataset from our earlier work, which included eCLIP datasets for 22 additional RBPs from two different cell lines, K562 and HepG2 [29]. The peak intensities on the RBP crosslinking sites were represented as the Irreproducible Discovery Rate (IDR) BED files [66], which were processed by the original authors. All the sequencing data were unified to the length of 100 nucleotides, and the samples with sequence identity over 80% were removed by using CD-HIT-EST module [67]. We kept the positive-to-negative ratio of 1:2 for each eCLIP-RBP, and the data partitioning outlined above were also complied to maintain an equitable model simulation scenario.

### 4.4 Human m6A modifications across nine cell lines

We downloaded single nucleotide resolution human m6A modification data including those obtained by different technologies and from nine cell lines [47]. The final dataset contains a total of 131,703 high-confidence m6A sites. We also derived a negative data set (total 108,573 nucleotides) by selecting non-positive adenosine nucleotides on the 3’UTR of the same transcript, using a non-overlapping window size of 41 nucleotides centered around the selected adenosine nucleotide. We ran CD-HIT-EST with a cut-off of 0.8 to remove potential redundancy between the positive and negative datasets, which produced a final 79,021 m6A sites and 849,005 non-m6A negative set (please refer to **Supplementary Fig. 7**). We noted that such 1:10 positive-negative ratio was also used in a previous study WHISTLE[49]. Following the balancing strategy described therein, we conducted random down-sampling for each cell-line by splitting the non-redundant negative samples into 10 subsets, each with 1:1 positive-to-negative ratio. Finally, we used 10-fold cross-validation to obtain independent test sets under unbiased partitions over all the subsets. The testing performances from the 10 independent sessions were averaged and reported as final results.

### 4.5 mRNA subcellular localization

RNA sub-cellular localization is regulated by specific cis-regulatory elements typically found in 5’ and 3’UTRs. We downloaded RNA localization data from the RNALocate database [55], as described in a previously published study [10]. Considering a single mRNA could have multiple annotated localizations including nucleus, exosome, cytosol, ribosome, membrane, and endoplasmic reticulum (ER), we merged the annotation of cytoplasm with cytosol since these two terms were ambiguous. A previous published method, DM3Loc [10], provided a benchmark dataset containing 17,023 mRNAs, which has removed sequence redundancy (80% cutoff) by running CD-HIT-EST [67]. The non-redundant dataset was further partitioned into training and validation set for 5-fold cross-validation, with each fold having similar distribution of subcellular localization categories. The five folds of test data outside the training process were used to evaluate the prediction performance. For each mRNA transcript, we followed the DM3Loc to take the first 4000 nts at the 5’ and the last 4000 nts at the 3’ and concatenated them into a single sequence; for those mRNA transcripts shorter than 8000 nts, we right-padded with zeros. In addition to benchmarking against the previously annotated dataset provided by DM3Loc, we also conducted an independent test on mRNAs that had annotations in the updated RNALocate v2.0 database but were not included in the DM3Loc dataset [54]. This novel dataset consisted of 2,615 mRNAs; the detailed sub-cellular locations could be found in **Supplementary Fig. 8**. After removing homologous sequences using CD-HIT-EST with 80% cutoff, a final set of 2,551 sequences was derived.

### 4.6 Tokenize mRNA sequences

An mRNA sequence is a string of nucleotides resembling human text; however, they lack explicit punctuation annotations to form discrete objects. Instead of considering each base as a single token, we merge consecutive nucleotides into *k-*mers since functional motifs usually span multiple nucleotides. Given an RNA sequence such as “AUGACA”, the 3-mers for this sequence consist of (AUG, UGA, GAC, ACA), or a sequence of two 5-mers (AUGAC, UGACA). We selected *k* from 3 to 6 to generate token representations of different length. The final *k*-mer vocabulary consists of a total of 4^k^ + 5 tokens including permutations of *k*-mer and 5 special encoding symbols. Specifically, classification token [CLS] represents global contextual semantics of the entire sequence; separation token [SEP] represents the end of each input sequence; unknown token [UNK] represents all *k*-mers that comprised at least one ambiguously sequenced character (e.g., “N”); masked token [MASK] represents the base fragment masked during self-supervised learning process; and padding token [PAD] represents padding shorter input sequences to the desired length [21].

### 4.7 Architecture of 3UTRBERT

3UTRBERT leverages deep representation learning to improve from task-specific approaches to a more generalized and broad range of post-transcriptional regulatory tasks. It uses pre-training on the unannotated 3’UTR data (total 20,362 sequences and 76,435,649 nucleotides) to learn contextual information among nucleotides, akin to learning grammar and syntax of human languages. These syntaxes or grammars in turn could reveal potential regulatory elements. The central component of 3UTRBERT architecture is the encoder part of a Transformer, which learns representations of the RNA universe via multi-head self-attention mechanism (please refer to **Fig. 1**). Firstly, the sequential segments of *k*-mer linguistic tokens *t* = < *t*_1_, *t*_2_,…,*t*_n_ > were fed into an Embedding layer (**E** ∈ ℝ^*n*×*d*^), which independently converts each token *t_i_* of vocabulary size *n* into a numerical vector with dimension *d* = 768. The positional embedding is then summed with token embedding as the final input of the Transformer encoder network.

We stacked 12 identical Transformer components as shared networks. Each layer contains a multi-head self-attention module and a position-wise fully connected feed-forward layer (FFN), with a residual connection around each sub-layer followed by a layer-wise normalization to greatly facilitate training. In particular, for the *k*-mer token embedding matrix **X**, the dot-product self-attention coefficient was calculated as the following:

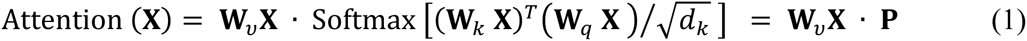

where W*_q_* ∈ ℝ*^d_q_×d^*, W_*k*_ ∈ ℝ*^d_k_×d^*, and W*_v_* ∈ ℝ*^d_v_×d^* represented the randomly initialized projection parameters associated with the *query*, *key*, and *value* respectively. For each single-head attention unit, we used *d_q_* = *d_k_* = *d_v_* = *d*, and the matrix **P** was designed to capture the contextual relevance of the input for a given token against the remaining tokens in the input sequence. Subsequently, the information flow of weighted average was delivered to the formula below:

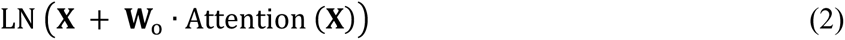

Here LN(·) denotes the layer-normalization operation and **W**_o_ ∈ ℝ^*d*×*d*^. To jointly model global dependencies among tokens in different representation subspaces, multi-head attention was independently performed to linearly project the *queries*, *ke*ys, and *values h* times (number of heads = 12) in parallel. This was followed by concatenation of the output hidden state from h different heads, resulting in the final values as formulated below:

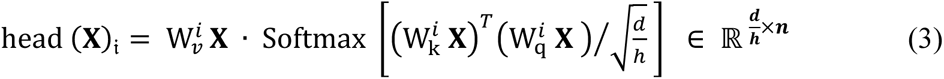

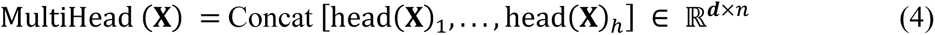

where the *query*, *key*, and *value* projection matrices 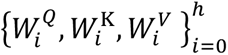 were 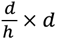 matrices. Since each head reduces the dimension to compute the context, the total calculation cost was close to that of a full-dimensional single-headed attention. The final multi-head attentional context layer then became:

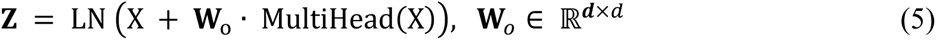

Following each attention layer, the discovered transcriptome contextual understanding passed through a fully connected feed-forward network to each position separately and identically, which consisted of a single linear transformation with size *h* = 3072 and a GeLU activation:

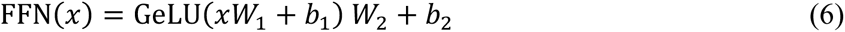

Compared with the CNN-based approaches which only captured the local context of nucleotide sequences, 3UTRBERT provides the flexibility to extract global dependency information from the entire input sequence. By performing self-attention mechanism on all the representations from the previous layer to calculate the hidden states, this model architecture could be easily parallelized; in addition, it could effectively overcome the gradient vanishing problem commonly encountered in RNN-based methods.

### 4.8 Self-supervised pre-training stage

To empower the language model to learn the syntax and semantics embedded in 3UTRs, we modified the Masked Language Model [MLM] pre-training task from the original deep representation learning implementation [18]. Given that mRNA 3’UTRs were not composed of multiple sentences, we discarded the Next Sentence Prediction (NSP) task. The essence of MLM is a self-supervised strategy of randomly masking a portion of an input 3’UTR sequence and then trains the network to predict their original values using the surrounding nucleotides. Specifically, 15% of the *k*-mers was randomly masked, and the model was iteratively optimized to restore the masked parts based on the rest. To reduce mismatches between pre-training and fine-tuning stages, 80% of the masked token was replaced by a special token [MASK], 10% was randomly substituted with a *k*-mer token from the vocabulary and another 10% remained unchanged from their original base. However, unlike human languages, the specific grammar of the *k*-mer representation introduced a problem in token masking as a shielded *k*-mer could be trivially inferred from immediate adjacent tokens. Taking a 3-mer sequence (AUG, UGA, GAC, ACA, CAG) as an example, “GAC” is the concatenation of “GA” in “UGA” and “C” in “ACA”. If we masked 15% of the *k*-mers randomly and independently, both the previous and the next *k*-mers of the masked *k*-mer would remain unmasked in most cases. To prevent the training task becoming simplified and thus affecting the learning of deep semantic relations, instead of masking each *k*-mer independently, we masked the *k*-mer for contiguous *k*-length spans, i.e., (AUG, MASK, MASK, MASK, CAG). After this, by building a classification layer upon the last hidden state of the Transformer, the output probability of each *k-*mer was calculated by a softmax function with cross-entropy loss:

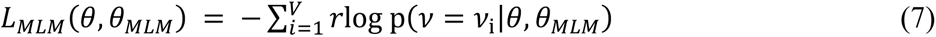

where *θ* and *θ_MLM_* respectively denoted the parameters of the 3UTRBERT pre-training model and the MLM task layer (multi-class classifier), and *V* equaled the number of tokens in the vocabulary. In addition, we experimented with different *k*-mer length in our model, as short *k*-mer took shorter time to train than longer *k*-mer but had less biological content; for example, 6-mer model takes 3 times more to train than 3-mer models. We set *k* as 3,4,5 and 6 to generate and evaluate four types of models. Each model followed the same architecture and parameter setting during pre-training for about 38 days using four NVIDIA Quadro RTX 6000 24G GPUs. Parallel training technique was employed on four GPUs to support the large batch size of 128. We optimized 3UTRBERT following the training tricks described in previous publications [15, 68] using AdamW (β_1_ = 0.9, β_2_ = 0.98, ε = 1*e*-6), which iterated for a total of 200k steps. The learning rate was linearly increased (i.e., warm-up) from 0 to 4e-4 in the first 10k steps and then linearly decreased to 0 after 200k steps with weight decay rate set as 0.01.

### 4.9 Supervised fine-tuning on several benchmarks

The agnostic pre-training step allowed 3UTRBERT to learn the syntax and grammar encoded in the 3’UTR of the mRNA transcripts; next we fine-tuned the model to perform several downstream tasks (**Fig. 1**). We showed that 3UTRBERT could transfer knowledge acquired in the pre-training step to understand specific follow-up post-transcriptional regulatory activities. Some of the downstream tasks took longer mRNA sequences as input, e.g., predicting mRNA sub-cellular localization using entire mRNA transcripts, whereas tasks such as predicting RBP binding sites or m6A sites took short sequence fragments as input. For long mRNA sequences, we froze the parameters inside 3UTRBERT and then fed the vector representations from the last hidden state (top right in **Fig. 1**) to the model of multi-label subcellular localization. It has been shown before that the representations on the feature-dimension with mean pooling could prevent fusion bias caused by excessive dimensionality [21]. In contrast, the short-sequence fragments were initialized from the pre-trained parameters and directly fine-tuned with task-specific data, which took much less time than pre-training. We provided the aggregate sequence representation from the special token [CLS] of final hidden state into an output layer for target classifications, this layer was usually composed of a single-layer feed-forward neural network and softmax classifier. When experimenting 3UTRBERT with different *k*-mers, we used identical hyperparameter settings and optimization tricks across all the applications, where AdamW fixed the weight decay (= 0.01) and used dropout (= 0.1) to alleviate the risk of overfitting. The learning rate was increased linearly to the peak value (= 5e-5) over the warm-up period and decayed to near 0. Among evaluating different *k*-mer sizes, we elected to report the final results for *k*-mer = 3 since it achieved the best performance; only slight fluctuations on performance across different *k*-mers was observed.

### 4.10 Visualization of 3UTRBERT and motif extraction

The rationale of 3UTRBERT is to identify nucleotide sequences that have high attention scores, which suggest that these nucleotides are potentially under functional constraints, e.g., binding sites for RBPs or m6A modification sites. Leveraging the self-attentive mechanism, 3UTRBERT was naturally applicable for mapping and deciphering regulatory regions in 3’UTRs. The special classification token [CLS] in first position of the input represents the aggregated representation from the last output layer [18]; the contribution score of each embedded *k*-mer token ±_*j*_ accumulated over all attention heads *H* in the context of “entire sequence” was calculated following the formula below:

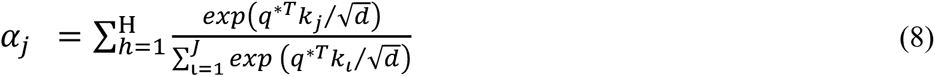

where *q*^∗^ denotes the query vector of [CLS] token with the dimension of *d*; *k_j_*_=_ denotes the key vector for the *j*-th *k*-mer token; and *J* was the number of tokens within input sequence. To convert the attention score from *k*-mer to individual nucleotides, for a particular nucleotide, we averaged the values of all the *k*-mers that the nucleotide was part of. After having identified sequence regions of high attention scores, we next evaluated whether they were enriched in RBP binding sites or m6A modification sites. The extracted attention weights from [CLS] to each token (±*_CLS_*, *j*) were used to identify high-attention regions in each positive sequence; sequence motif candidates were identified with 7-mer length. We then used hypergeometric test to calculate the statistical significance of the enrichment of these identified motifs and conducted Benjamini-Hochberg multi-testing correction with adjusted p-value < 0.005. Finally, we performed pairwise alignment to merge similar motif candidates, the resulting instances with the highest numbers were converted into a position-weight matrix (PWM) format. These motifs were then compared with the experimentally validated motifs stored in ATtRACT database [31] using TOMTOM algorithm [32].

## 5. DECLARATIONS

### Ethics approval and consent to participate

Not Applicable.

### Consent for publication

Not Applicable.

### Availability of data and materials

All datasets in this study were derived from published resources and can be generated following protocols described in **Methods**. Source code, demo data, and tutorial is available on GitHub at: https://github.com/yangyn533/3UTRBERT; the complete dataset can be downloaded from https://figshare.com/articles/dataset/3UTRBERT_dataset_availability/22845644.

### Competing interests

The authors declare that they have no competing interests.

### Funding

Z.Z. acknowledges support from the Nature Science and Engineering Research Council of Canada (NSERC, RGPIN-2017-06743). X.L. acknowledges support from National Natural Science Foundation of China (NSFC, Grant No. 62076109) and the Jilin Province Outstanding Young Scientist Program (Grant No. 20230508098RC), and the Fundamental Research Funds for the Central Universities awarded to Jilin University.

### Authors’ contributions

Y.Y., X.L. and Z.Z. conceived the study. Y.Y., G.L., K.P., and W.C. developed the methods and conducted the analysis. Y.Y., X.L. and Z.Z. wrote the manuscript, and all authors revised it. All authors read and approved the final version of the manuscript.

## Acknowledgements

We thank Dr. Hongli Ma for her help with the RBP-RNA binding data analysis.

## Notes

https://github.com/yangyn533/3UTRBERT

### Competing Interest Statement

The authors have declared no competing interest.

