## Supplementary Info for "Deciphering 3’ UTR mediated gene regulation using interpretable deep representation learning"

### Supplementary Note 1: Definition of the evaluation metrics

| Short name | Full name | Formula |
|---|---|---|
| ACC | Accuracy | $\mathrm{ACC} = \frac{\mathrm{TP+TN}}{\mathrm{TP+FP+FN+TN}}$ |
| AUROC | Area Under the Receiver Operating Characteristic | ROC AUC is the area under the curve<br>where x is false positive rate(FPR)<br>and the y is true positive rate(TPR). |
| AUPRC | Area Under the Precision-Recall Curve | PR AUC is the area under the curve<br>where x is recall and y is precision. |
| MCC | Matthews correlation coefficient | $\mathrm{MCC} = \frac{TP*TN-FP*FN}{\sqrt{(TP+FP)*(TP+FN)*(TN+FP)*(TN+FN)}}$ |
| F1 | F1-measure<br>(harmonic mean of precision and sensitivity) | $\mathrm{F1} = \frac{2*PPR*TPR}{PPR+TPR} = \frac{2*TP}{2*TP+FP+FN}$ |
| Precision | Precision | $\mathrm{Precision} = \frac{TP}{TP+FP}$ |
| Recall | Recall | $\mathrm{Recall} = \frac{TP}{TP+FN}$ |
| TPR | True positive rate(sensitivity) | $\mathrm{TPR} = \frac{TP}{TP+FN}$ |
| FPR | False positive rate | $\mathrm{FPR} = \frac{FP}{TN+FP}$ |
| PPR | Predicted positive rate(precision) | $\mathrm{PPR} = \frac{TP}{TP+FP}$ |
| TP | True positive | Number of correctly predicted crosslink/modification sites. |
| TN | True negative | Number of correctly predicted<br>non-crosslink/non-modification residues. |
| FP | False positive | Number of non-crosslink/non-modification residues<br>incorrectly predicted as crosslink/modification. |
| FN | False negative | Number of crosslink/modification residues incorrectly<br>predicted as non-crosslink/non-modification. |

For sequence labelling tasks of binary classification, the predictions were generated from the propensities such that nucleotides with propensities greater than a given threshold are identified as crosslinked/modified, and otherwise they are identified as non-crosslinked/ non-modified. We evaluated the predictive performance of the binary identifications with the four metrics and thresholds as 0.5: Accuracy, F1, and MCC (Matthews correlation coefficient). Accuracy showed us how comfortable the model was with detecting the positive and negative classes. It was computed by the sum of True Positives and True Negatives divided by the total population. F1 ranged between 0 and 1 where higher value denoted more accurate prediction. MCC ranged between –1 and 1, where –1 represented an inverted prediction (all predictions were flipped compared to the experimental values), 0 denoted a random result and 1 denoted a perfect prediction. The area under the receiver operating characteristic (AUROC) curve to evaluate discriminate quality of the propensities. The AUROC curve was a relation between true positive rates (TPRs) and false-positive rates (FPRs) that was calculated by thresholding the propensities where the thresholds were the set of all unique propensities produced by a given predictor. ROC-AUC ranged between 0.5 (equivalent to a random identification) and 1 (perfect identification). To support multi-label classification, the estimator was wrapped in a OneVsRestClassifier to produce binary comparisons for each class (e.g. the positive case is the class and the negative case is any other class). The precision-recall curve (AUPRC) showed the tradeoff between precision and recall for all classes. The AUPRC curve provided a more accurate assessment of the model's performance than metrics such as accuracy or F1 score, which might be biased towards the majority class.

### Supplementary Note 2: RNA-protein docking and molecular dynamics simulation

In this study, the 3D structures of the filtered RNA sequences were modelled with the DeepFoldRNA software (1), a stand-alone program obtained from the official release at https://github.com/robpearc/DeepFoldRNA. Commencing with an RNA sequence, the program generated a homology-based alignment of numerous sequences sourced from diverse databases (as of March 2023). Subsequently, the software employed deep neural networks to predict spatial restraints, which were transformed into negative log-likelihood potentials. Finally, L-BFGS folding simulations were executed to produce a full-length structure model. Since no experimentally determined structure was available for RBM15, we resorted to computational methods for structure prediction. Specifically, we obtained the protein structure of RBM15 from AlphaFoldDB, a publicly database of protein structures generated by the AlphaFold algorithm (2). We retrieved the protein structure from AlphaFoldDB (entry Q96T37) and selected the RNA Recognition Motif (RRM) to build the molecular dynamics (MD) system. Crystal structure for SND1 was retrieved from PDB entry 5M9O, and further utilized to perform the analysis. The ClusPro (3) web server was subsequently employed to model the RNA-protein docking, and the populated binding pattern was sent to the MD simulation for evaluating the stability and dynamics. After that, the Charmm-gui webserver was applied to construct the MD systems (4–6). The RNA-protein complex was solvated in a water box with a 15 Å buffer around it and the system was neutralized with K+ and Cl- ions at 0.15M concentration. The systems were equilibrated and energy-minimized for 10,000 steps in the NPT ensemble. We then stabilized the temperature at T = 30°C with the Nosé-Hoover method (7) and the coupled pressure with the Parrinello–Rahman method (8). A switch distance of 10-Å and a cutoff distance of 12-Å for non-bonded interactions were applied. Bond lengths were further constrained with hydrogen atoms using LINCS (9) with a 2-fs time step, and the systems varied in the number of atoms from 216,000 to 332,000. We finally performed energy minimization and production MD simulations with GROMACS (2021.6) (10), using CHARMM36m force field (11) and TIP3P water model (12).

### Supplementary Figure 1

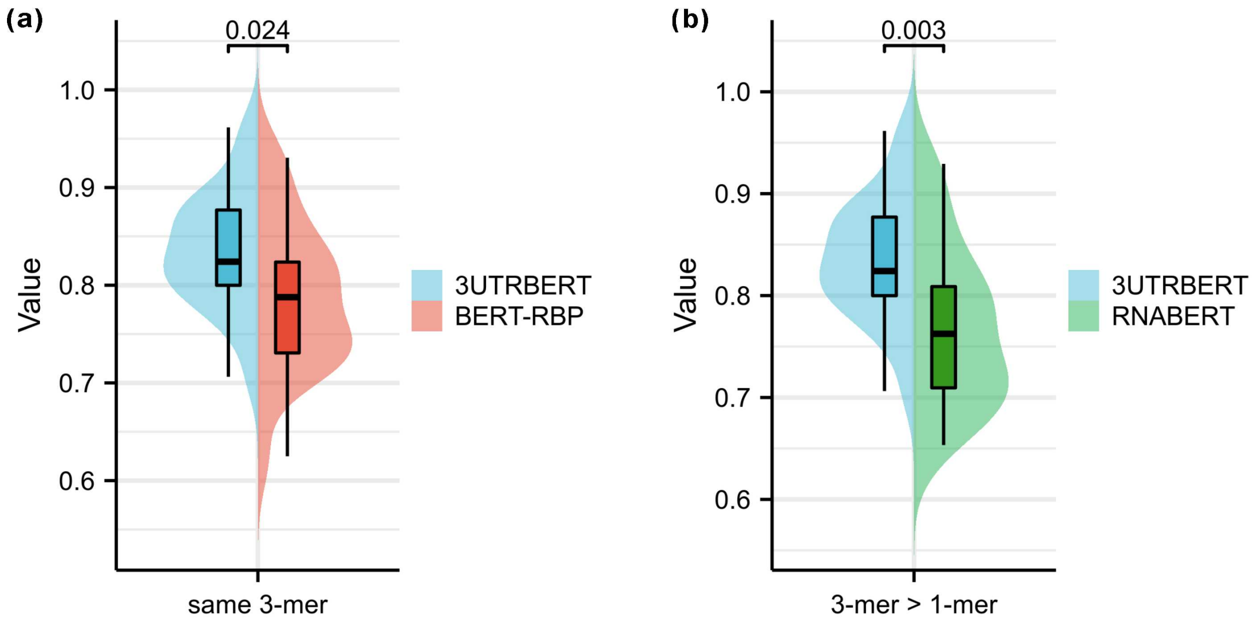


**Supplementary Fig. 1.** Comparison results of Transformer architecture-based approaches for 22 RBPs under eCLIP protocol. (a) At the same token length, 3UTRBERT injected with prior knowledge of the regulatory regions was superior to BERT_RBP, which was directly fine-tuned on the human reference genome and lacked the pre-training stage, in identifying RNA-protein cross-linking stable sites. (b) Since none of the regulatory motifs in the non-coding region occurred at single resolution, RNABERT thus suffered from a performance bottleneck caused by the loss of local contextual information. More importantly, RNABERT had difficulty in modelling the relationship between UTR and target protein expressions based on the implicit representation learned by non-coding RNAs.

### Supplementary Figure 2

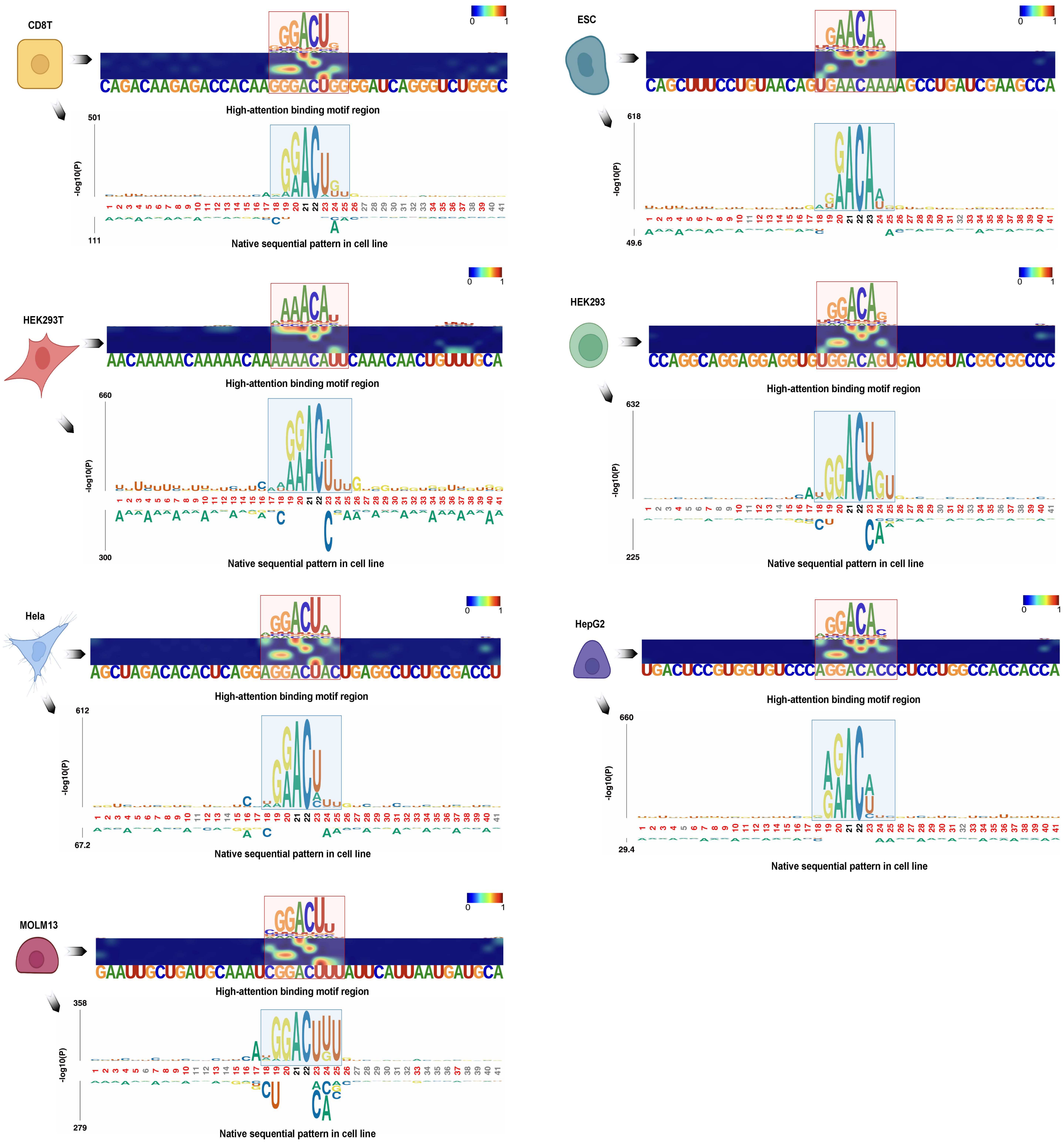


**Supplementary Fig. 2.** 3UTRBERT explored the consensus patterns of sequential information across various cell lines, thus equipping superior generalizability to dynamically identify cellular epitranscriptomic modifications.

### Supplementary Figure 3

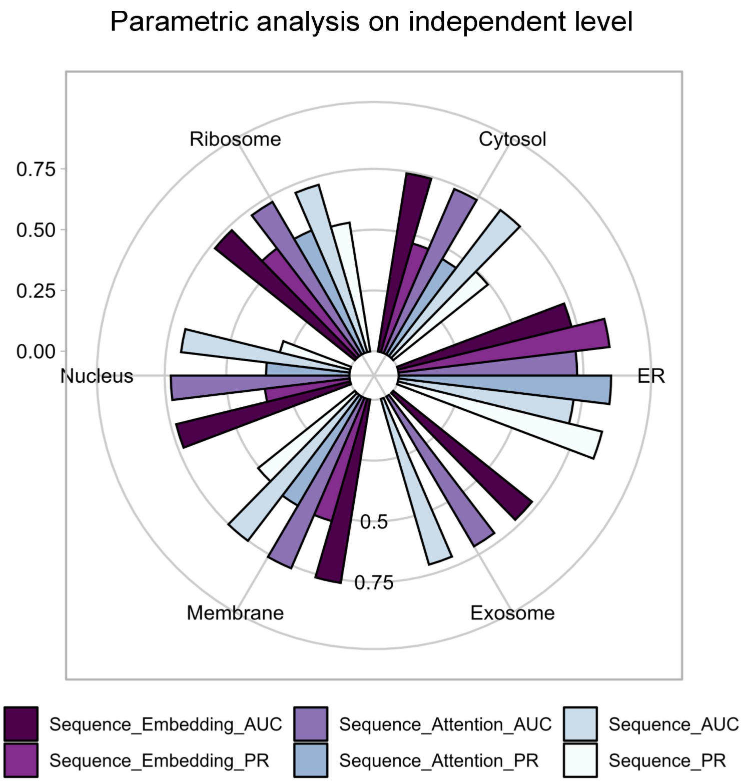


**Supplementary Fig. 3.** Circular histogram plots the performance on identical model architecture of different feature schemes for multi-label mRNA subcellular localization prediction.

### Supplementary Figure 4

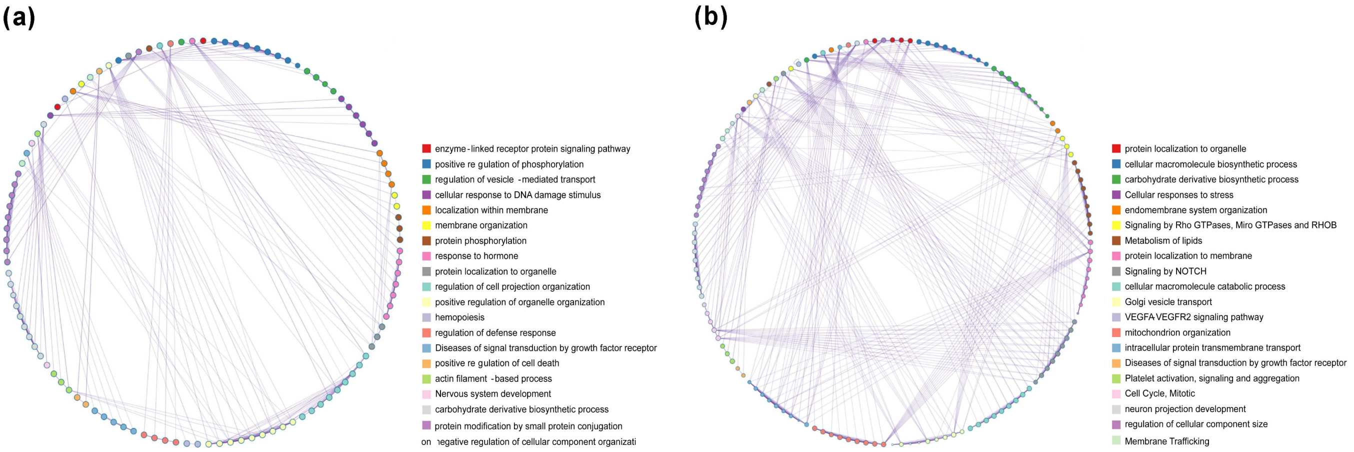


**Supplementary Fig. 4.** The Gene Ontology Network respectively displayed the relationships contained in Endoplasmic Reticulum (right) and the sets composed of nucleus, exosome, cytosol, and membrane (left) on the independent test set. The contextual semantics embedding generated by 3UTRBERT yielded better performance for ER compartment with more abundant background information.

### Supplementary Figure 5

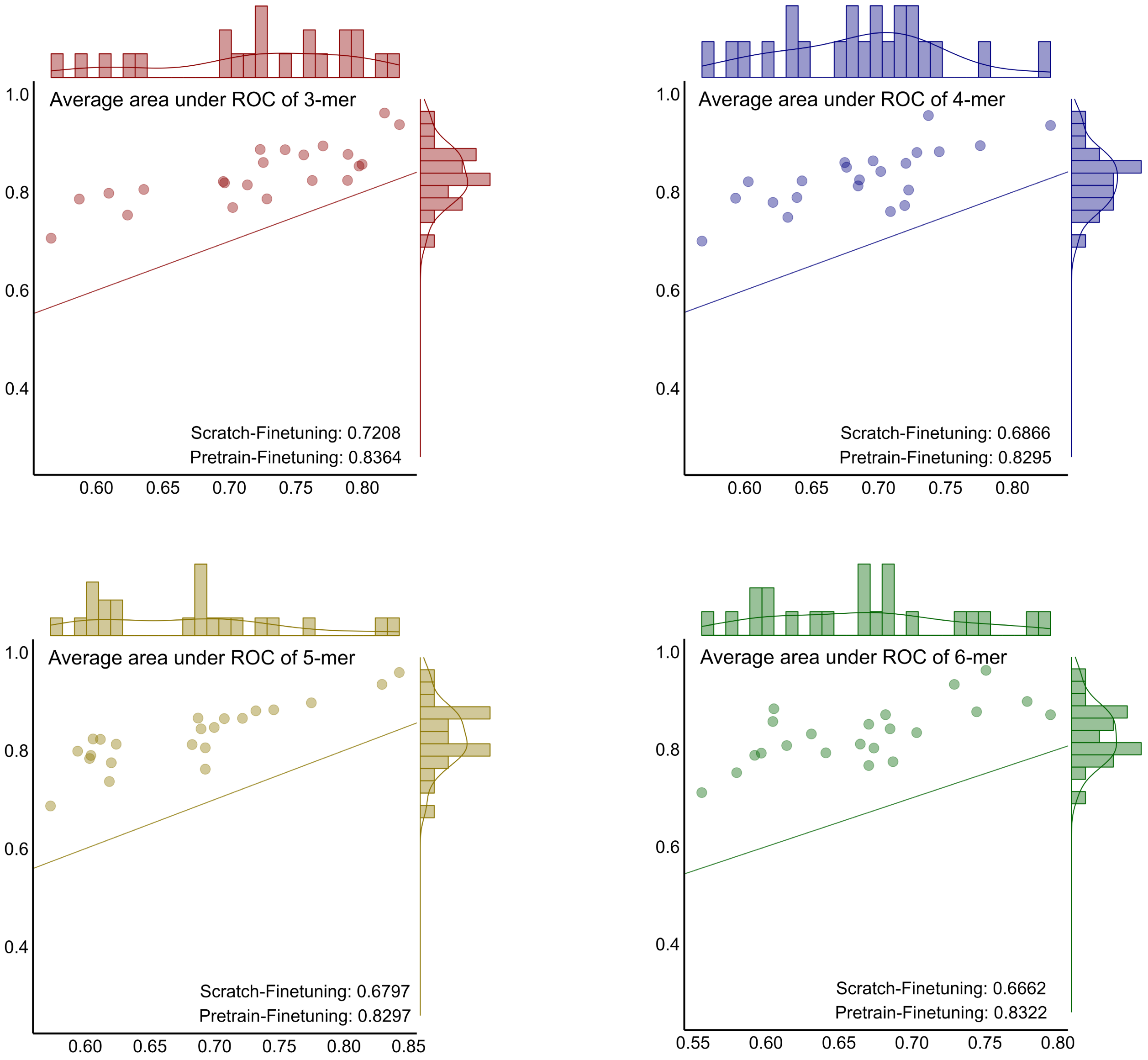


**Supplementary Fig. 5.** Performance comparison on 22 RBPs between the stage of scratch-finetuning and pretrain-finetuning in terms of AUCs to demonstrate that the task-agnostic self-supervised learning approach captured generalized and transferable understanding from regulatory regions compared with random initialization information.

### Supplementary Figure 6

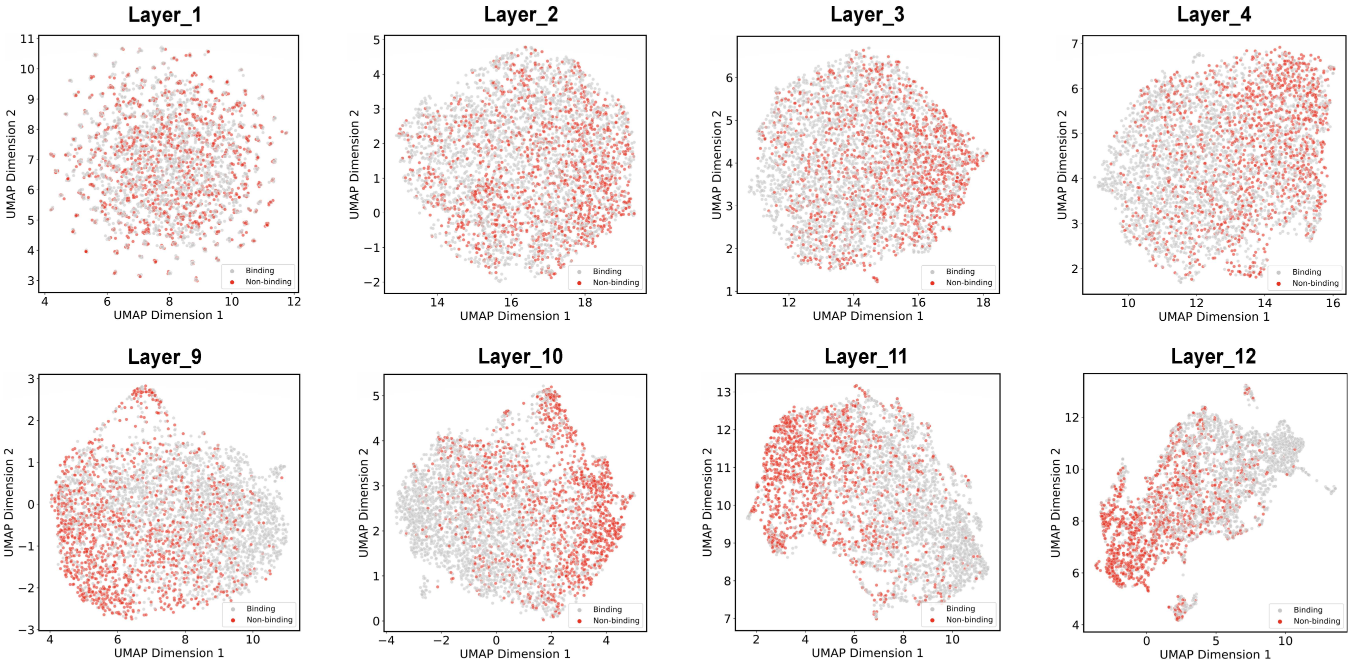


**Supplementary Fig. 6.** A two-dimensional projection of the embedding vectors generated by 3UTRBERT using UMAP to visualize the clustering effect of each layer on RNA sequences; linkage sites (positive samples) were annotated in colors while non-linkage sites (negative samples) with grey color. As moving up the layers in language model, the representations become increasingly contextualized, which make them more effective for classification tasks.

### Supplementary Figure 7

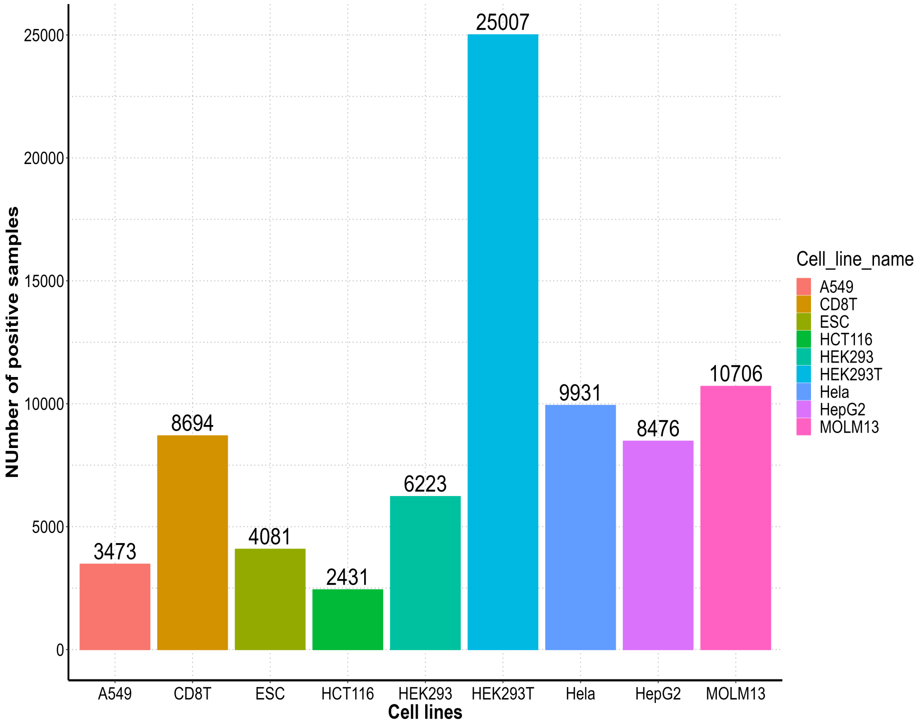


**Supplementary Fig. 7.** The human epitranscriptomic modifications distribution across different nine cell lines.

### Supplementary Figure 8

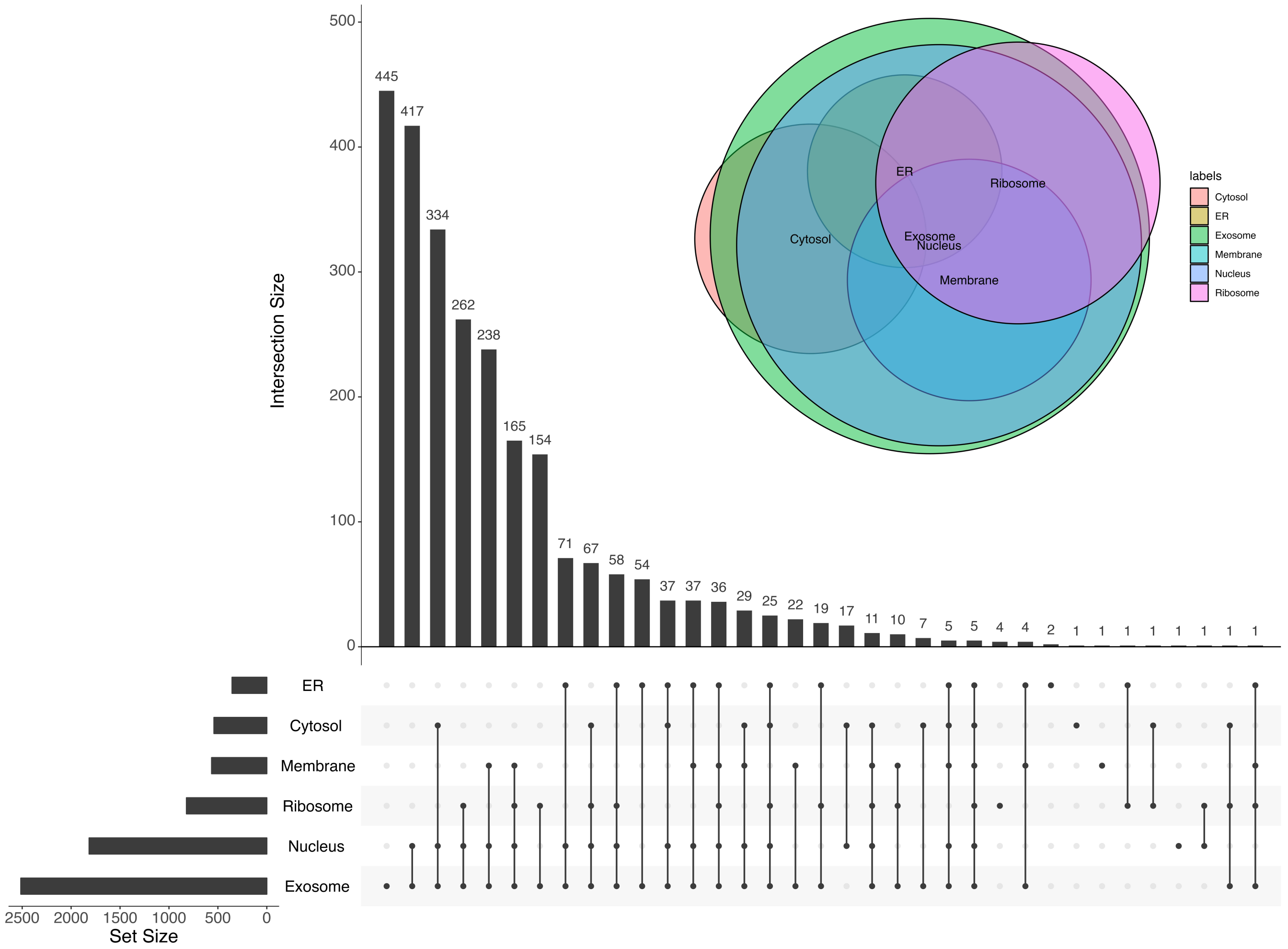


**Supplementary Fig. 8.** The venn upset showed the detailed number of mRNA sequences in each intersection group on independent test set. Upper bar plot presented the number of mRNAs in each intersection group, while the left bar plot indicated the total number of sequences for each subcompartment. And the bottom dots displayed the components of each group.

**Supplementary Table. 1.** Performance evaluation in terms of average AUCs, F1-score, MCC and ACC with Std for 3UTRBERT, BERT_RBP, RNABERT, GraphProt2, RPI_Net, DeepCLIP and iDeepE on 22 eCLIP protocols.

| Dataset_22_**AUCs** | 3UTRBERT | BERT_RBP | RNABERT | GraphProt2 | RPI_Net | DeepCLIP | iDeepE |
|---|---|---|---|---|---|---|---|
| AKAP1_HepG2 | **0.887**±0.004 | 0.860±0.001 | 0.841±0.001 | 0.844±0.007 | 0.767±0.006 | 0.749±0.005 | 0.744±0.008 |
| BCLAF1_HepG2 | **0.877**±0.002 | 0.807±0.003 | 0.805±0.004 | 0.787±0.002 | 0.722±0.009 | 0.741±0.001 | 0.718±0.006 |
| DDX3X_HepG2 | **0.938**±0.002 | 0.931±0.002 | 0.929±0.001 | 0.923±0.010 | 0.851±0.004 | 0.593±0.008 | 0.849±0.005 |
| DDX3X_K562 | 0.895±0.005 | 0.886±0.001 | 0.877±0.003 | 0.876±0.004 | 0.799±0.009 | **0.920**±0.004 | 0.805±0.004 |
| DDX24_K562 | 0.798±0.005 | 0.734±0.007 | 0.676±0.016 | 0.703±0.006 | 0.610±0.014 | **0.829**±0.011 | 0.624±0.010 |
| FAM120A_K562 | **0.861**±0.002 | 0.825±0.005 | 0.802±0.028 | 0.820±0.010 | 0.724±0.010 | 0.725±0.009 | 0.741±0.009 |
| G3BP1_HepG2 | **0.786**±0.006 | 0.724±0.008 | 0.653±0.044 | 0.668±0.010 | 0.611±0.015 | 0.558±0.006 | 0.606±0.005 |
| GRWD1_HepG2 | **0.806**±0.003 | 0.720±0.006 | 0.698±0.003 | 0.688±0.005 | 0.628±0.014 | 0.597±0.009 | 0.626±0.008 |
| IGF2BP1_K562 | **0.853**±0.003 | 0.807±0.003 | 0.788±0.004 | 0.784±0.012 | 0.697±0.017 | 0.747±0.008 | 0.724±0.005 |
| LARP4_HepG2 | **0.769**±0.004 | 0.728±0.006 | 0.708±0.008 | 0.709±0.012 | 0.625±0.009 | 0.652±0.010 | 0.626±0.004 |
| LIN28B_K562 | **0.706**±0.004 | 0.625±0.019 | 0.696±0.004 | 0.610±0.011 | 0.539±0.023 | 0.561±0.007 | 0.558±0.004 |
| PABPC4_K562 | **0.787**±0.003 | 0.730±0.008 | 0.713±0.003 | 0.725±0.012 | 0.633±0.010 | 0.658±0.011 | 0.634±0.010 |
| PPIG_HepG2 | **0.876**±0.003 | 0.821±0.005 | 0.780±0.005 | 0.760±0.003 | 0.691±0.032 | 0.689±0.008 | 0.685±0.007 |
| PUM2_K562 | **0.962**±0.001 | 0.924±0.002 | 0.916±0.002 | 0.941±0.004 | 0.877±0.011 | 0.837±0.008 | 0.855±0.007 |
| RBM15_K562 | **0.824**±0.002 | 0.771±0.004 | 0.745±0.007 | 0.721±0.007 | 0.692±0.013 | 0.685±0.008 | 0.687±0.004 |
| RPS3_HepG2 | **0.824**±0.004 | 0.805±0.003 | 0.784±0.003 | 0.752±0.008 | 0.698±0.012 | 0.744±0.011 | 0.677±0.014 |
| SND1_HepG2 | **0.815**±0.002 | 0.764±0.005 | 0.734±0.004 | 0.714±0.008 | 0.655±0.009 | 0.635±0.003 | 0.652±0.014 |
| UCHL5_K562 | **0.822**±0.004 | 0.749±0.005 | 0.736±0.001 | 0.726±0.004 | 0.657±0.009 | 0.660±0.003 | 0.659±0.003 |
| UPF1_HepG2 | **0.887**±0.002 | 0.865±0.002 | 0.860±0.002 | 0.843±0.006 | 0.772±0.008 | 0.773±0.006 | 0.770±0.003 |
| UPF1_K562 | **0.857**±0.003 | 0.820±0.002 | 0.810±0.002 | 0.799±0.009 | 0.723±0.015 | 0.725±0.006 | 0.721±0.005 |
| YBX3_K562 | **0.753**±0.004 | 0.702±0.004 | 0.668±0.006 | 0.651±0.010 | 0.688±0.023 | 0.559±0.007 | 0.591±0.004 |
| ZNF622_K562 | **0.819**±0.006 | 0.738±0.005 | 0.713±0.007 | 0.731±0.005 | 0.653±0.006 | 0.650±0.006 | 0.662±0.005 |
| Avg. ± Std. | **0.836**±0.003 | 0.788±0.005 | 0.770±0.007 | 0.762±0.007 | 0.696±0.013 | 0.695±0.007 | 0.692±0.007 |

| Dataset_22_**F1** | 3UTRBERT | BERT_RBP | RNABERT | GraphProt2 | RPI_Net | DeepCLIP | iDeepE |
|---|---|---|---|---|---|---|---|
| AKAP1_HepG2 | **0.799**±0.005 | 0.691±0.008 | 0.652±0.009 | 0.660±0.023 | 0.691±0.007 | 0.450±0.022 | 0.655±0.012 |
| BCLAF1_HepG2 | **0.785**±0.002 | 0.537±0.013 | 0.565±0.011 | 0.550±0.017 | 0.626±0.015 | 0.445±0.026 | 0.619±0.009 |
| DDX3X_HepG2 | **0.862**±0.004 | 0.806±0.006 | 0.804±0.005 | 0.797±0.010 | 0.803±0.003 | 0.100±0.028 | 0.802±0.005 |
| DDX3X_K562 | **0.834**±0.003 | 0.757±0.003 | 0.753±0.007 | 0.728±0.019 | 0.735±0.012 | 0.783±0.011 | 0.745±0.005 |
| DDX24_K562 | **0.708**±0.012 | 0.488±0.018 | 0.317±0.040 | 0.432±0.016 | 0.438±0.031 | 0.643±0.018 | 0.469±0.017 |
| FAM120A_K562 | **0.765**±0.007 | 0.619±0.022 | 0.597±0.050 | 0.598±0.015 | 0.632±0.017 | 0.401±0.016 | 0.658±0.012 |
| G3BP1_HepG2 | **0.692**±0.004 | 0.486±0.016 | 0.240±0.140 | 0.373±0.018 | 0.442±0.037 | 0.078±0.024 | 0.422±0.013 |
| GRWD1_HepG2 | **0.717**±0.004 | 0.452±0.013 | 0.378±0.040 | 0.375±0.023 | 0.473±0.034 | 0.072±0.039 | 0.463±0.018 |
| IGF2BP1_K562 | **0.760**±0.007 | 0.539±0.032 | 0.510±0.031 | 0.546±0.027 | 0.590±0.030 | 0.451±0.012 | 0.632±0.008 |
| LARP4_HepG2 | **0.691**±0.004 | 0.412±0.021 | 0.398±0.034 | 0.396±0.028 | 0.443±0.024 | 0.183±0.032 | 0.451±0.008 |
| LIN28B_K562 | **0.625**±0.008 | 0.206±0.100 | 0.476±0.020 | 0.200±0.040 | 0.198±0.111 | 0.026±0.016 | 0.311±0.015 |
| PABPC4_K562 | **0.708**±0.004 | 0.461±0.014 | 0.442±0.035 | 0.449±0.027 | 0.465±0.022 | 0.165±0.021 | 0.463±0.022 |
| PPIG_HepG2 | **0.784**±0.003 | 0.625±0.007 | 0.550±0.019 | 0.514±0.023 | 0.572±0.060 | 0.291±0.058 | 0.568±0.013 |
| PUM2_K562 | **0.905**±0.002 | 0.787±0.013 | 0.780±0.004 | 0.812±0.013 | 0.840±0.012 | 0.659±0.013 | 0.810±0.006 |
| RBM15_K562 | **0.731**±0.003 | 0.522±0.028 | 0.460±0.037 | 0.440±0.035 | 0.577±0.024 | 0.330±0.031 | 0.567±0.008 |
| RPS3_HepG2 | **0.724**±0.003 | 0.590±0.010 | 0.534±0.018 | 0.462±0.051 | 0.587±0.021 | 0.447±0.034 | 0.555±0.026 |
| SND1_HepG2 | **0.715**±0.008 | 0.505±0.058 | 0.426±0.049 | 0.355±0.044 | 0.512±0.019 | 0.087±0.027 | 0.501±0.027 |
| UCHL5_K562 | **0.744**±0.003 | 0.492±0.024 | 0.512±0.040 | 0.475±0.021 | 0.515±0.020 | 0.333±0.023 | 0.516±0.006 |
| UPF1_HepG2 | **0.798**±0.003 | 0.700±0.005 | 0.695±0.011 | 0.636±0.011 | 0.695±0.009 | 0.503±0.017 | 0.695±0.004 |
| UPF1_K562 | **0.765**±0.003 | 0.626±0.004 | 0.607±0.008 | 0.585±0.018 | 0.631±0.025 | 0.388±0.037 | 0.627±0.008 |
| YBX3_K562 | **0.666**±0.005 | 0.348±0.044 | 0.206±0.064 | 0.222±0.043 | 0.354±0.069 | 0.001±0.001 | 0.367±0.012 |
| ZNF622_K562 | **0.737**±0.003 | 0.478±0.027 | 0.407±0.037 | 0.473±0.023 | 0.504±0.014 | 0.290±0.036 | 0.526±0.012 |
| Avg. ± Std. | **0.751**±0.005 | 0.551±0.022 | 0.514±0.032 | 0.503±0.025 | 0.560±0.028 | 0.324±0.025 | 0.565±0.012 |

| Dataset_22_**MCC** | 3UTRBERT | BERT_RBP | RNABERT | GraphProt2 | RPI_Net | DeepCLIP | iDeepE |
|---|---|---|---|---|---|---|---|
| AKAP1_HepG2 | **0.599**±0.010 | 0.541±0.006 | 0.493±0.009 | 0.512±0.019 | 0.542±0.010 | 0.315±0.012 | 0.509±0.014 |
| BCLAF1_HepG2 | **0.571**±0.004 | 0.418±0.013 | 0.404±0.005 | 0.373±0.007 | 0.472±0.011 | 0.250±0.017 | 0.461±0.007 |
| DDX3X_HepG2 | 0.724±0.008 | 0.713±0.006 | 0.711±0.008 | 0.701±0.015 | 0.707±0.002 | **0.773**±0.013 | 0.706±0.006 |
| DDX3X_K562 | 0.670±0.006 | 0.652±0.005 | 0.644±0.008 | 0.619±0.021 | 0.616±0.013 | **0.681**±0.013 | 0.636±0.005 |
| DDX24_K562 | 0.417±0.021 | 0.311±0.019 | 0.201±0.025 | 0.269±0.006 | 0.255±0.021 | **0.508**±0.026 | 0.272±0.020 |
| FAM120A_K562 | **0.531**±0.013 | 0.459±0.012 | 0.419±0.054 | 0.430±0.020 | 0.467±0.008 | 0.255±0.018 | 0.498±0.021 |
| G3BP1_HepG2 | **0.385**±0.007 | 0.290±0.006 | 0.144±0.061 | 0.207±0.028 | 0.251±0.024 | 0.148±0.009 | 0.252±0.008 |
| GRWD1_HepG2 | **0.434**±0.008 | 0.285±0.013 | 0.229±0.014 | 0.227±0.011 | 0.286±0.013 | 0.257±0.023 | 0.286±0.012 |
| IGF2BP1_K562 | **0.523**±0.013 | 0.390±0.022 | 0.347±0.019 | 0.369±0.028 | 0.405±0.018 | 0.275±0.013 | 0.464±0.006 |
| LARP4_HepG2 | **0.382**±0.008 | 0.287±0.013 | 0.265±0.018 | 0.263±0.015 | 0.311±0.008 | 0.143±0.022 | 0.303±0.009 |
| LIN28B_K562 | **0.259**±0.014 | 0.116±0.029 | 0.243±0.009 | 0.103±0.019 | 0.132±0.046 | 0.128±0.023 | 0.155±0.010 |
| PABPC4_K562 | **0.418**±0.006 | 0.327±0.013 | 0.295±0.013 | 0.306±0.026 | 0.316±0.011 | 0.168±0.016 | 0.323±0.016 |
| PPIG_HepG2 | **0.568**±0.006 | 0.459±0.007 | 0.368±0.006 | 0.344±0.006 | 0.415±0.047 | 0.158±0.025 | 0.396±0.010 |
| PUM2_K562 | **0.811**±0.004 | 0.679±0.016 | 0.661±0.007 | 0.722±0.018 | 0.760±0.016 | 0.513±0.015 | 0.715±0.005 |
| RBM15_K562 | **0.464**±0.005 | 0.367±0.011 | 0.310±0.025 | 0.273±0.030 | 0.412±0.015 | 0.191±0.011 | 0.410±0.005 |
| RPS3_HepG2 | **0.449**±0.006 | 0.405±0.006 | 0.349±0.013 | 0.280±0.030 | 0.408±0.014 | 0.278±0.019 | 0.375±0.015 |
| SND1_HepG2 | **0.430**±0.015 | 0.338±0.027 | 0.258±0.018 | 0.214±0.015 | 0.333±0.011 | 0.333±0.008 | 0.334±0.020 |
| UCHL5_K562 | **0.490**±0.005 | 0.359±0.013 | 0.301±0.027 | 0.322±0.017 | 0.361±0.013 | 0.218±0.016 | 0.370±0.008 |
| UPF1_HepG2 | **0.596**±0.005 | 0.557±0.011 | 0.552±0.010 | 0.497±0.008 | 0.561±0.008 | 0.365±0.020 | 0.555±0.006 |
| UPF1_K562 | **0.530**±0.006 | 0.455±0.004 | 0.438±0.004 | 0.415±0.016 | 0.457±0.006 | 0.278±0.016 | 0.462±0.005 |
| YBX3_K562 | **0.340**±0.008 | 0.243±0.018 | 0.147±0.022 | 0.147±0.006 | 0.230±0.028 | 0.305±0.020 | 0.236±0.003 |
| ZNF622_K562 | **0.475**±0.007 | 0.341±0.019 | 0.271±0.019 | 0.327±0.016 | 0.361±0.006 | 0.195±0.011 | 0.368±0.006 |
| Avg. ± Std. | **0.503**±0.009 | 0.409±0.013 | 0.366±0.018 | 0.360±0.017 | 0.412±0.016 | 0.306±0.016 | 0.413±0.010 |

| Dataset_22_**ACC** | 3UTRBERT | BERT_RBP | RNABERT | GraphProt2 | RPI_Net | DeepCLIP | iDeepE |
|---|---|---|---|---|---|---|---|
| AKAP1_HepG2 | **0.822**±0.004 | 0.798±0.002 | 0.780±0.003 | 0.789±0.007 | 0.798±0.005 | 0.722±0.003 | 0.790±0.005 |
| BCLAF1_HepG2 | **0.812**±0.002 | 0.759±0.005 | 0.750±0.003 | 0.733±0.002 | 0.772±0.003 | 0.689±0.003 | 0.770±0.002 |
| DDX3X_HepG2 | **0.877**±0.003 | 0.874±0.003 | 0.873±0.003 | 0.869±0.007 | 0.870±0.001 | 0.671±0.001 | 0.870±0.002 |
| DDX3X_K562 | 0.856±0.003 | 0.849±0.002 | 0.845±0.003 | 0.839±0.008 | 0.834±0.005 | **0.860**±0.005 | 0.843±0.002 |
| DDX24_K562 | 0.746±0.008 | 0.718±0.007 | 0.694±0.006 | 0.704±0.002 | 0.694±0.005 | **0.794**±0.011 | 0.700±0.007 |
| FAM120A_K562 | **0.789**±0.006 | 0.764±0.004 | 0.746±0.020 | 0.757±0.009 | 0.767±0.003 | 0.700±0.006 | 0.778±0.010 |
| G3BP1_HepG2 | **0.735**±0.005 | 0.706±0.002 | 0.681±0.005 | 0.685±0.011 | 0.692±0.009 | 0.669±0.002 | 0.701±0.003 |
| GRWD1_HepG2 | **0.748**±0.004 | 0.712±0.005 | 0.698±0.001 | 0.694±0.003 | 0.703±0.003 | 0.671±0.004 | 0.710±0.003 |
| IGF2BP1_K562 | **0.791**±0.004 | 0.745±0.007 | 0.727±0.006 | 0.735±0.010 | 0.741±0.003 | 0.704±0.006 | 0.766±0.006 |
| LARP4_HepG2 | **0.731**±0.005 | 0.720±0.003 | 0.712±0.004 | 0.719±0.006 | 0.730±0.003 | 0.697±0.004 | 0.722±0.004 |
| LIN28B_K562 | **0.694**±0.004 | 0.676±0.004 | 0.675±0.005 | 0.680±0.003 | 0.685±0.004 | 0.682±0.003 | 0.675±0.005 |
| PABPC4_K562 | **0.753**±0.003 | 0.733±0.004 | 0.721±0.003 | 0.728±0.010 | 0.729±0.003 | 0.701±0.003 | 0.731±0.004 |
| PPIG_HepG2 | **0.810**±0.001 | 0.767±0.003 | 0.732±0.002 | 0.729±0.004 | 0.755±0.014 | 0.679±0.002 | 0.745±0.004 |
| PUM2_K562 | **0.914**±0.002 | 0.856±0.006 | 0.844±0.005 | 0.878±0.008 | 0.893±0.007 | 0.790±0.006 | 0.872±0.002 |
| RBM15_K562 | **0.771**±0.002 | 0.740±0.001 | 0.723±0.006 | 0.707±0.009 | 0.752±0.005 | 0.686±0.003 | 0.755±0.001 |
| RPS3_HepG2 | **0.758**±0.005 | 0.743±0.002 | 0.726±0.006 | 0.708±0.008 | 0.747±0.004 | 0.711±0.004 | 0.736±0.003 |
| SND1_HepG2 | **0.760**±0.005 | 0.734±0.003 | 0.712±0.003 | 0.703±0.007 | 0.728±0.008 | 0.682±0.004 | 0.736±0.005 |
| UCHL5_K562 | **0.780**±0.002 | 0.737±0.005 | 0.699±0.002 | 0.724±0.006 | 0.736±0.004 | 0.695±0.003 | 0.740±0.003 |
| UPF1_HepG2 | **0.820**±0.002 | 0.806±0.003 | 0.804±0.003 | 0.787±0.003 | 0.811±0.004 | 0.740±0.006 | 0.806±0.003 |
| UPF1_K562 | **0.789**±0.003 | 0.763±0.002 | 0.758±0.000 | 0.749±0.006 | 0.758±0.010 | 0.708±0.006 | 0.767±0.001 |
| YBX3_K562 | **0.740**±0.004 | 0.726±0.002 | 0.709±0.002 | 0.711±0.008 | 0.721±0.007 | 0.709±0.011 | 0.720±0.001 |
| ZNF622_K562 | **0.771**±0.004 | 0.730±0.006 | 0.706±0.005 | 0.726±0.009 | 0.738±0.002 | 0.690±0.004 | 0.736±0.003 |
| Avg. ± Std. | **0.785**±0.004 | 0.757±0.004 | 0.742±0.004 | 0.743±0.007 | 0.757±0.005 | 0.711±0.004 | 0.758±0.004 |

**Supplementary Table. 2.** Performance evaluation in terms of average AUCs, F1-score, Mcc and Acc with Std for 3UTRBERT, BERT_RBP, RNABERT, GraphProt2, RPI_Net, DeepCLIP and iDeepE on 31 datasets covering CLIP/HITS-CLIP, iCLIP and PAR-CLIP from hg19.

| Dataset_31_**AUCs** | 3UTRBERT | BERT_RBP | RNABERT | GraphProt2 | RPI_Net | DeepCLIP | iDeepE |
|---|---|---|---|---|---|---|---|
| AGO2_PARCLIP | 0.684±0.012 | 0.720±0.014 | 0.649±0.006 | 0.712±0.022 | **0.745**±0.010 | 0.572±0.016 | 0.654±0.013 |
| AGO2-M_PARCLIP | **0.671**±0.010 | 0.633±0.015 | 0.627±0.005 | 0.600±0.017 | 0.633±0.008 | 0.579±0.021 | 0.513±0.012 |
| AGO1234_PARCLIP | **0.813**±0.008 | 0.609±0.015 | 0.792±0.004 | 0.657±0.012 | 0.669±0.008 | 0.518±0.009 | 0.631±0.028 |
| Binding_1_HITSCLIP | **0.893**±0.005 | 0.888±0.008 | 0.685±0.002 | 0.841±0.016 | 0.638±0.011 | 0.558±0.020 | 0.641±0.004 |
| Binding_2_HITSCLIP | **0.894**±0.004 | 0.871±0.004 | 0.675±0.003 | 0.845±0.011 | 0.652±0.019 | 0.578±0.011 | 0.624±0.010 |
| eIF4AIII_1_CLIPSEQ | **0.965**±0.002 | 0.947±0.004 | 0.813±0.002 | 0.930±0.006 | 0.810±0.021 | 0.841±0.008 | 0.823±0.014 |
| eIF4AIII_2_CLIPSEQ | **0.964**±0.003 | 0.955±0.005 | 0.865±0.001 | 0.938±0.008 | 0.826±0.013 | 0.846±0.011 | 0.806±0.013 |
| ELVAL1-1_PARCLIP | 0.911±0.004 | **0.919**±0.004 | 0.898±0.001 | 0.883±0.018 | 0.775±0.014 | 0.804±0.009 | 0.763±0.020 |
| ELVAL1-2__PARCLIP | **0.925**±0.003 | **0.925**±0.004 | 0.898±0.001 | 0.906±0.013 | 0.840±0.019 | 0.475±0.015 | 0.813±0.019 |
| ELVAL1-A_PARCLIP | 0.892±0.004 | **0.894**±0.006 | 0.836±0.001 | 0.850±0.007 | 0.789±0.017 | 0.752±0.015 | 0.761±0.018 |
| ELVAL1-M_PARCLIP | 0.639±0.013 | 0.628±0.014 | **0.852**±0.008 | 0.582±0.020 | 0.662±0.020 | 0.762±0.013 | 0.660±0.039 |
| EWSR1_PARCLIP | 0.893±0.006 | **0.900**±0.007 | 0.876±0.007 | 0.810±0.009 | 0.774±0.005 | 0.603±0.023 | 0.755±0.011 |
| FUS_PARCLIP | 0.930±0.008 | **0.932**±0.005 | 0.890±0.002 | 0.823±0.015 | 0.850±0.020 | 0.620±0.010 | 0.815±0.010 |
| hnRNPC-1_ICLIP | 0.959±0.002 | **0.960**±0.002 | 0.931±0.002 | 0.903±0.011 | 0.857±0.012 | 0.696±0.033 | 0.840±0.020 |
| hnRNPC-2_ICLIP | **0.978**±0.001 | 0.976±0.002 | 0.960±0.002 | 0.937±0.007 | 0.884±0.015 | 0.626±0.020 | 0.877±0.003 |
| hnRNPL-1_ICLIP | 0.695±0.014 | 0.654±0.013 | **0.849**±0.002 | 0.663±0.016 | 0.663±0.011 | 0.832±0.018 | 0.596±0.014 |
| hnRNPL-2_ICLIP | 0.665±0.011 | 0.613±0.011 | **0.931**±0.002 | 0.651±0.012 | 0.616±0.008 | 0.827±0.013 | 0.642±0.019 |
| HnRNPL-L_ICLIP | 0.675±0.009 | 0.658±0.008 | **0.849**±0.005 | 0.647±0.016 | 0.509±0.006 | 0.626±0.038 | 0.578±0.017 |
| IGF2BP1-3_PARCLIP | **0.805**±0.003 | 0.707±0.019 | 0.746±0.005 | 0.676±0.031 | 0.737±0.005 | 0.723±0.023 | 0.777±0.006 |
| MOV10_PARCLIP | 0.819±0.006 | 0.836±0.004 | **0.849**±0.003 | 0.792±0.005 | 0.844±0.004 | 0.669±0.019 | 0.601±0.016 |
| mut-FUS_PARCLIP | 0.937±0.003 | **0.940**±0.002 | 0.872±0.006 | 0.846±0.012 | 0.890±0.013 | 0.793±0.013 | 0.821±0.015 |
| NSUN2_ICLIP | **0.823**±0.006 | 0.807±0.009 | 0.781±0.008 | 0.762±0.010 | 0.593±0.028 | 0.771±0.008 | 0.691±0.011 |
| PUM2_PARCLIP | 0.967±0.001 | **0.974**±0.002 | 0.890±0.001 | 0.931±0.010 | 0.815±0.020 | 0.782±0.010 | 0.839±0.009 |
| QKI_PARCLIP | **0.956**±0.005 | **0.956**±0.004 | 0.928±0.001 | 0.936±0.011 | 0.902±0.011 | 0.791±0.013 | 0.885±0.020 |
| SFRS1_CLIPSEQ | **0.910**±0.003 | 0.891±0.005 | 0.827±0.005 | 0.856±0.013 | 0.695±0.015 | 0.752±0.023 | 0.706±0.015 |
| TAF15_PARCLIP | **0.950**±0.005 | 0.936±0.013 | 0.882±0.011 | 0.853±0.006 | 0.894±0.008 | 0.810±0.004 | 0.861±0.012 |
| TDP-43_ICLIP | 0.914±0.009 | **0.918**±0.004 | 0.850±0.002 | 0.882±0.008 | 0.773±0.026 | 0.778±0.009 | 0.776±0.011 |
| TIA1_ICLIP | **0.933**±0.003 | 0.918±0.004 | 0.896±0.005 | 0.870±0.012 | 0.770±0.009 | 0.818±0.012 | 0.789±0.020 |
| TIAL1_ICLIP | **0.883**±0.003 | 0.878±0.006 | 0.869±0.008 | 0.836±0.011 | 0.752±0.011 | 0.602±0.012 | 0.731±0.008 |
| U2AF65_ICLIP | **0.958**±0.004 | **0.958**±0.005 | 0.924±0.001 | 0.857±0.009 | 0.807±0.005 | 0.621±0.021 | 0.825±0.023 |
| Y2AF65_ICLIP | **0.930**±0.003 | 0.922±0.005 | 0.882±0.006 | 0.857±0.011 | 0.785±0.015 | 0.763±0.019 | 0.766±0.008 |
| Avg. ± Std. | **0.865**±0.006 | 0.849±0.007 | 0.841±0.004 | 0.811±0.012 | 0.756±0.013 | 0.703±0.016 | 0.737±0.015 |

| Dataset_31_**F1** | 3UTRBERT | BERT_RBP | RNABERT | GraphProt2 | RPI_Net | DeepCLIP | iDeepE |
|---|---|---|---|---|---|---|---|
| AGO2_PARCLIP | **0.573**±0.028 | 0.149±0.028 | 0.518±0.038 | 0.217±0.066 | 0.099±0.020 | 0.165±0.013 | 0.518±0.029 |
| AGO2-M_PARCLIP | **0.546**±0.007 | 0.053±0.019 | 0.524±0.033 | 0.064±0.056 | 0.072±0.042 | 0.191±0.023 | 0.098±0.028 |
| AGO1234_PARCLIP | 0.651±0.012 | 0.056±0.065 | **0.684**±0.020 | 0.514±0.031 | 0.106±0.022 | 0.098±0.020 | 0.444±0.071 |
| Binding_1_HITSCLIP | **0.779**±0.011 | 0.594±0.029 | 0.342±0.040 | 0.524±0.019 | 0.430±0.025 | 0.144±0.021 | 0.435±0.010 |
| Binding_2_HITSCLIP | **0.772**±0.007 | 0.528±0.021 | 0.274±0.023 | 0.521±0.048 | 0.453±0.031 | 0.171±0.020 | 0.398±0.023 |
| eIF4AIII_1_CLIPSEQ | **0.896**±0.005 | 0.792±0.013 | 0.344±0.064 | 0.722±0.022 | 0.710±0.021 | 0.509±0.012 | 0.743±0.019 |
| eIF4AIII_2_CLIPSEQ | **0.892**±0.005 | 0.770±0.010 | 0.535±0.039 | 0.722±0.028 | 0.732±0.017 | 0.518±0.024 | 0.705±0.011 |
| ELVAL1-1_PARCLIP | **0.782**±0.014 | 0.633±0.019 | 0.594±0.031 | 0.559±0.073 | 0.641±0.015 | 0.460±0.018 | 0.628±0.022 |
| ELVAL1-2_PARCLIP | **0.839**±0.005 | 0.735±0.017 | 0.653±0.013 | 0.608±0.016 | 0.734±0.015 | 0.089±0.019 | 0.714±0.024 |
| ELVAL1-A_PARCLIP | **0.796**±0.008 | 0.580±0.060 | 0.548±0.017 | 0.536±0.024 | 0.666±0.018 | 0.356±0.033 | 0.634±0.024 |
| ELVAL1-M_PARCLIP | **0.558**±0.016 | 0.102±0.044 | 0.331±0.147 | 0.055±0.030 | 0.132±0.037 | 0.387±0.025 | 0.509±0.087 |
| EWSR1_PARCLIP | **0.796**±0.013 | 0.656±0.051 | 0.484±0.071 | 0.409±0.043 | 0.656±0.010 | 0.239±0.029 | 0.634±0.021 |
| FUS_PARCLIP | **0.844**±0.008 | 0.722±0.025 | 0.622±0.042 | 0.403±0.049 | 0.752±0.018 | 0.194±0.026 | 0.717±0.009 |
| hnRNPC-1_ICLIP | **0.877**±0.006 | 0.801±0.010 | 0.696±0.028 | 0.646±0.021 | 0.773±0.012 | 0.248±0.050 | 0.762±0.025 |
| hnRNPC-2_ICLIP | **0.901**±0.010 | 0.834±0.008 | 0.789±0.007 | 0.710±0.034 | 0.818±0.011 | 0.220±0.019 | 0.823±0.007 |
| hnRNPL-1_ICLIP | **0.599**±0.013 | 0.121±0.021 | 0.365±0.041 | 0.172±0.049 | 0.158±0.024 | 0.485±0.032 | 0.336±0.032 |
| hnRNPL-2_ICLIP | 0.547±0.030 | 0.135±0.021 | **0.702**±0.014 | 0.092±0.042 | 0.061±0.020 | 0.502±0.036 | 0.480±0.041 |
| HnRNPL-L_ICLIP | **0.592**±0.018 | 0.156±0.018 | 0.292±0.111 | 0.178±0.041 | 0.039±0.026 | 0.108±0.010 | 0.289±0.041 |
| IGF2BP1-3_PARCLIP | 0.329±0.016 | 0.094±0.029 | 0.592±0.015 | 0.154±0.035 | 0.120±0.035 | 0.296±0.020 | **0.702**±0.009 |
| MOV10_PARCLIP | **0.653**±0.027 | 0.324±0.070 | 0.501±0.025 | 0.391±0.028 | 0.414±0.039 | 0.301±0.034 | 0.349±0.036 |
| mut-FUS_PARCLIP | **0.851**±0.014 | 0.725±0.046 | 0.607±0.029 | 0.474±0.052 | 0.790±0.005 | 0.440±0.030 | 0.730±0.017 |
| NSUN2_ICLIP | **0.677**±0.003 | 0.459±0.039 | 0.393±0.080 | 0.304±0.036 | 0.322±0.066 | 0.412±0.018 | 0.523±0.019 |
| PUM2_PARCLIP | **0.898**±0.006 | 0.821±0.009 | 0.554±0.048 | 0.705±0.026 | 0.711±0.022 | 0.416±0.034 | 0.762±0.008 |
| QKI_PARCLIP | **0.900**±0.006 | 0.857±0.026 | 0.701±0.015 | 0.770±0.015 | 0.847±0.011 | 0.424±0.034 | 0.829±0.021 |
| SFRS1_CLIPSEQ | **0.806**±0.005 | 0.629±0.010 | 0.430±0.010 | 0.542±0.035 | 0.530±0.022 | 0.396±0.043 | 0.556±0.026 |
| TAF15_PARCLIP | **0.871**±0.011 | 0.669±0.071 | 0.580±0.062 | 0.449±0.035 | 0.801±0.006 | 0.263±0.101 | 0.782±0.018 |
| TDP-43_ICLIP | **0.823**±0.008 | 0.725±0.008 | 0.497±0.051 | 0.680±0.018 | 0.688±0.034 | 0.421±0.010 | 0.688±0.018 |
| TIA1_ICLIP | **0.817**±0.005 | 0.619±0.035 | 0.656±0.036 | 0.519±0.027 | 0.638±0.014 | 0.459±0.021 | 0.673±0.023 |
| TIAL1_ICLIP | **0.752**±0.007 | 0.567±0.029 | 0.566±0.030 | 0.526±0.033 | 0.603±0.014 | 0.210±0.017 | 0.589±0.016 |
| U2AF65_ICLIP | **0.869**±0.008 | 0.774±0.030 | 0.716±0.005 | 0.528±0.032 | 0.692±0.007 | 0.240±0.011 | 0.740±0.031 |
| Y2AF65_ICLIP | **0.826**±0.006 | 0.656±0.034 | 0.622±0.034 | 0.537±0.020 | 0.656±0.022 | 0.384±0.032 | 0.641±0.008 |
| Avg. ± Std. | **0.752**±0.011 | 0.527±0.029 | 0.539±0.039 | 0.459±0.035 | 0.511±0.022 | 0.314±0.027 | 0.594±0.025 |

| Dataset_31_**MCC** | 3UTRBERT | BERT_RBP | RNABERT | GraphProt2 | RPI_Net | DeepCLIP | iDeepE |
|---|---|---|---|---|---|---|---|
| AGO2_PARCLIP | **0.504**±0.043 | 0.155±0.013 | 0.121±0.033 | 0.168±0.046 | 0.117±0.021 | 0.023±0.018 | 0.330±0.011 |
| AGO2-M_PARCLIP | **0.461**±0.021 | 0.101±0.042 | 0.127±0.031 | 0.033±0.060 | 0.121±0.035 | 0.049±0.027 | 0.058±0.052 |
| AGO1234_PARCLIP | **0.572**±0.021 | 0.044±0.051 | 0.382±0.020 | 0.356±0.038 | 0.070±0.043 | 0.139±0.021 | 0.340±0.022 |
| Binding_1_HITSCLIP | **0.560**±0.020 | 0.520±0.028 | 0.314±0.023 | 0.451±0.019 | 0.406±0.026 | 0.016±0.030 | 0.392±0.013 |
| Binding_2_HITSCLIP | **0.544**±0.014 | 0.455±0.019 | 0.245±0.008 | 0.444±0.037 | 0.404±0.003 | 0.400±0.020 | 0.363±0.029 |
| eIF4AIII_1_CLIPSEQ | **0.793**±0.010 | 0.741±0.015 | 0.271±0.032 | 0.663±0.022 | 0.645±0.021 | 0.401±0.018 | 0.689±0.021 |
| eIF4AIII_2_CLIPSEQ | **0.785**±0.010 | 0.717±0.011 | 0.454±0.031 | 0.664±0.027 | 0.669±0.020 | 0.409±0.030 | 0.638±0.010 |
| ELVAL1-1_PARCLIP | 0.564±0.028 | **0.566**±0.013 | 0.503±0.027 | 0.474±0.075 | 0.555±0.016 | 0.340±0.020 | 0.540±0.023 |
| ELVAL1-2__PARCLIP | 0.677±0.011 | **0.682**±0.019 | 0.571±0.013 | 0.539±0.009 | 0.668±0.019 | 0.528±0.013 | 0.648±0.027 |
| ELVAL1-A_PARCLIP | **0.592**±0.016 | 0.531±0.052 | 0.447±0.015 | 0.452±0.021 | 0.586±0.019 | 0.225±0.031 | 0.555±0.023 |
| ELVAL1-M_PARCLIP | 0.165±0.013 | 0.132±0.049 | 0.302±0.104 | 0.066±0.037 | 0.153±0.031 | 0.259±0.030 | **0.345**±0.040 |
| EWSR1_PARCLIP | **0.593**±0.025 | **0.593**±0.050 | 0.429±0.056 | 0.319±0.033 | 0.581±0.014 | 0.096±0.036 | 0.562±0.027 |
| FUS_PARCLIP | 0.689±0.016 | 0.660±0.028 | 0.544±0.029 | 0.322±0.044 | **0.691**±0.021 | 0.047±0.024 | 0.651±0.009 |
| hnRNPC-1_ICLIP | **0.755**±0.013 | 0.753±0.012 | 0.624±0.022 | 0.576±0.026 | 0.718±0.015 | 0.100±0.057 | 0.709±0.027 |
| hnRNPC-2_ICLIP | **0.804**±0.020 | 0.794±0.009 | 0.738±0.010 | 0.648±0.033 | 0.774±0.012 | 0.082±0.024 | 0.784±0.010 |
| hnRNPL-1_ICLIP | 0.216±0.019 | 0.057±0.037 | 0.323±0.024 | 0.160±0.024 | 0.059±0.030 | **0.371**±0.043 | 0.272±0.036 |
| hnRNPL-2_ICLIP | 0.133±0.060 | 0.060±0.034 | **0.631**±0.015 | 0.086±0.034 | 0.052±0.036 | 0.394±0.043 | 0.323±0.021 |
| HnRNPL-L_ICLIP | 0.215±0.035 | 0.162±0.028 | **0.266**±0.069 | 0.166±0.018 | 0.080±0.040 | 0.012±0.043 | 0.235±0.041 |
| IGF2BP1-3_PARCLIP | 0.266±0.024 | 0.106±0.007 | 0.238±0.014 | 0.101±0.032 | 0.134±0.014 | 0.153±0.028 | **0.566**±0.014 |
| MOV10_PARCLIP | 0.322±0.044 | 0.291±0.043 | **0.414**±0.021 | 0.304±0.026 | 0.341±0.030 | 0.170±0.038 | 0.264±0.023 |
| mut-FUS_PARCLIP | 0.703±0.027 | 0.673±0.045 | 0.523±0.021 | 0.375±0.047 | **0.738**±0.007 | 0.320±0.034 | 0.669±0.018 |
| NSUN2_ICLIP | **0.362**±0.005 | 0.360±0.026 | 0.329±0.046 | 0.236±0.037 | 0.259±0.030 | 0.283±0.016 | 0.435±0.021 |
| PUM2_PARCLIP | **0.796**±0.012 | 0.780±0.010 | 0.492±0.029 | 0.638±0.032 | 0.644±0.022 | 0.490±0.033 | 0.708±0.008 |
| QKI_PARCLIP | 0.800±0.011 | **0.821**±0.033 | 0.646±0.010 | 0.717±0.020 | 0.810±0.013 | 0.530±0.036 | 0.790±0.024 |
| SFRS1_CLIPSEQ | **0.615**±0.010 | 0.555±0.011 | 0.342±0.010 | 0.478±0.041 | 0.448±0.025 | 0.115±0.039 | 0.488±0.030 |
| TAF15_PARCLIP | 0.744±0.024 | 0.616±0.067 | 0.491±0.061 | 0.375±0.025 | **0.751**±0.008 | 0.272±0.058 | 0.728±0.023 |
| TDP-43_ICLIP | 0.660±0.012 | **0.670**±0.009 | 0.451±0.023 | 0.631±0.011 | 0.656±0.022 | 0.297±0.014 | 0.644±0.021 |
| TIA1_ICLIP | **0.637**±0.008 | 0.565±0.024 | 0.571±0.039 | 0.427±0.028 | 0.553±0.017 | 0.341±0.027 | 0.596±0.023 |
| TIAL1_ICLIP | 0.506±0.015 | **0.510**±0.022 | 0.463±0.027 | 0.435±0.038 | 0.506±0.016 | 0.263±0.021 | 0.504±0.023 |
| U2AF65_ICLIP | **0.739**±0.016 | 0.729±0.030 | 0.642±0.007 | 0.434±0.036 | 0.617±0.011 | 0.298±0.029 | 0.682±0.034 |
| Y2AF65_ICLIP | **0.653**±0.011 | 0.591±0.031 | 0.526±0.037 | 0.453±0.027 | 0.572±0.027 | 0.253±0.036 | 0.562±0.006 |
| Avg. ± Std. | **0.562**±0.020 | 0.483±0.028 | 0.433±0.029 | 0.393±0.032 | 0.464±0.021 | 0.248±0.030 | 0.518±0.023 |

| Dataset_31_**ACC** | 3UTRBERT | BERT_RBP | RNABERT | GraphProt2 | RPI_Net | DeepCLIP | iDeepE |
|---|---|---|---|---|---|---|---|
| AGO2_PARCLIP | **0.856**±0.003 | 0.803±0.002 | 0.794±0.008 | 0.796±0.006 | 0.801±0.002 | 0.736±0.009 | 0.720±0.004 |
| AGO2-M_PARCLIP | **0.841**±0.006 | 0.802±0.003 | 0.792±0.006 | 0.791±0.005 | 0.804±0.002 | 0.740±0.008 | 0.786±0.010 |
| AGO1234_PARCLIP | **0.867**±0.006 | 0.795±0.003 | 0.822±0.006 | 0.734±0.007 | 0.786±0.016 | 0.733±0.006 | 0.737±0.006 |
| Binding_1_HITSCLIP | **0.864**±0.005 | 0.859±0.006 | 0.821±0.003 | 0.844±0.004 | 0.841±0.005 | 0.745±0.011 | 0.835±0.003 |
| Binding_2_HITSCLIP | **0.856**±0.004 | 0.844±0.005 | 0.810±0.003 | 0.840±0.007 | 0.837±0.006 | 0.746±0.007 | 0.830±0.007 |
| eIF4AIII_1_CLIPSEQ | **0.932**±0.003 | 0.918±0.004 | 0.804±0.004 | 0.897±0.006 | 0.891±0.005 | 0.818±0.009 | 0.905±0.007 |
| eIF4AIII_2_CLIPSEQ | **0.930**±0.004 | 0.912±0.003 | 0.841±0.005 | 0.897±0.007 | 0.898±0.005 | 0.820±0.010 | 0.888±0.001 |
| ELVAL1-1_PARCLIP | **0.860**±0.006 | 0.871±0.003 | 0.846±0.002 | 0.845±0.017 | 0.859±0.008 | 0.800±0.005 | 0.856±0.004 |
| ELVAL1-2__PARCLIP | **0.898**±0.004 | 0.904±0.005 | 0.866±0.003 | 0.865±0.002 | 0.892±0.005 | 0.748±0.010 | 0.891±0.007 |
| ELVAL1-A_PARCLIP | **0.872**±0.004 | 0.866±0.012 | 0.831±0.004 | 0.839±0.009 | 0.870±0.003 | 0.773±0.007 | 0.864±0.005 |
| ELVAL1-M_PARCLIP | 0.787±0.009 | 0.803±0.003 | 0.821±0.013 | **0.798**±0.003 | 0.804±0.004 | 0.781±0.009 | 0.733±0.006 |
| EWSR1_PARCLIP | 0.871±0.008 | **0.879**±0.012 | 0.842±0.010 | 0.810±0.007 | 0.873±0.005 | 0.746±0.010 | 0.870±0.008 |
| FUS_PARCLIP | **0.902**±0.005 | 0.895±0.008 | 0.860±0.006 | 0.814±0.008 | 0.900±0.007 | 0.735±0.012 | 0.891±0.003 |
| hnRNPC-1_ICLIP | **0.922**±0.003 | **0.922**±0.004 | 0.878±0.004 | 0.873±0.008 | 0.911±0.005 | 0.741±0.017 | 0.910±0.007 |
| hnRNPC-2_ICLIP | **0.938**±0.006 | 0.935±0.003 | 0.917±0.004 | 0.892±0.009 | 0.929±0.003 | 0.747±0.009 | 0.933±0.003 |
| hnRNPL-1_ICLIP | 0.781±0.012 | 0.777±0.014 | **0.821**±0.004 | 0.800±0.004 | 0.764±0.017 | 0.809±0.016 | 0.809±0.008 |
| hnRNPL-2_ICLIP | 0.779±0.011 | 0.773±0.018 | **0.883**±0.006 | 0.795±0.005 | 0.792±0.010 | 0.817±0.013 | 0.733±0.003 |
| HnRNPL-L_ICLIP | 0.791±0.008 | 0.803±0.004 | **0.813**±0.007 | 0.801±0.005 | 0.803±0.002 | 0.799±0.002 | 0.805±0.007 |
| IGF2BP1-3_PARCLIP | 0.808±0.006 | 0.799±0.003 | 0.803±0.004 | 0.785±0.011 | 0.801±0.004 | 0.754±0.011 | **0.811**±0.007 |
| MOV10_PARCLIP | 0.808±0.006 | 0.817±0.005 | **0.831**±0.005 | 0.806±0.014 | 0.820±0.007 | 0.765±0.008 | 0.801±0.006 |
| mut-FUS_PARCLIP | 0.904±0.006 | 0.902±0.011 | 0.854±0.001 | 0.818±0.009 | **0.910**±0.004 | 0.797±0.009 | 0.898±0.005 |
| NSUN2_ICLIP | 0.813±0.002 | 0.813±0.007 | 0.818±0.005 | 0.802±0.007 | 0.806±0.006 | 0.784±0.007 | **0.836**±0.005 |
| PUM2_PARCLIP | **0.934**±0.004 | 0.931±0.003 | 0.854±0.002 | 0.888±0.010 | 0.889±0.006 | 0.786±0.010 | 0.910±0.002 |
| QKI_PARCLIP | 0.935±0.004 | **0.943**±0.011 | 0.894±0.001 | 0.912±0.007 | 0.940±0.004 | 0.789±0.007 | 0.934±0.007 |
| SFRS1_CLIPSEQ | **0.881**±0.004 | 0.867±0.004 | 0.816±0.003 | 0.851±0.011 | 0.840±0.010 | 0.801±0.003 | 0.853±0.008 |
| TAF15_PARCLIP | 0.915±0.006 | 0.886±0.016 | 0.845±0.011 | 0.826±0.003 | **0.916**±0.005 | 0.818±0.007 | 0.914±0.008 |
| TDP-43_ICLIP | 0.899±0.003 | **0.900**±0.003 | 0.847±0.005 | 0.890±0.004 | 0.899±0.006 | 0.790±0.007 | 0.895±0.006 |
| TIA1_ICLIP | **0.888**±0.003 | 0.873±0.005 | 0.857±0.008 | 0.832±0.007 | 0.861±0.006 | 0.801±0.009 | 0.874±0.005 |
| TIAL1_ICLIP | 0.847±0.005 | **0.860**±0.004 | 0.830±0.008 | 0.834±0.013 | 0.844±0.008 | 0.737±0.011 | 0.852±0.008 |
| U2AF65_ICLIP | 0.916±0.004 | **0.917**±0.008 | 0.877±0.007 | 0.832±0.011 | 0.879±0.005 | 0.745±0.023 | 0.902±0.009 |
| Y2AF65_ICLIP | **0.888**±0.003 | 0.878±0.007 | 0.842±0.008 | 0.841±0.009 | 0.864±0.009 | 0.778±0.009 | 0.866±0.001 |
| Avg. ± Std. | **0.871**±0.005 | 0.863±0.006 | 0.850±0.005 | 0.834±0.008 | 0.856±0.006 | 0.773±0.009 | 0.840±0.006 |

**Supplementary Table. 3.** Performance evaluation in terms of average AUCs, F1-score, MCC and ACC with Std for 3UTRBERT, DeepM6ASeq, iMRM, WHISTLE, and SCRAMP on human m6A modifications across nine cell lines.

| **AUCs** | 3UTRBERT | DeepM6ASeq | WHISTLE | iMRM | SCRAMP |
|---|---|---|---|---|---|
| A549 | **0.979**±0.003 | 0.941±0.008 | 0.879±0.002 | 0.918±0.006 | 0.912±0.003 |
| CD8T | **0.979**±0.003 | 0.959±0.003 | 0.861±0.001 | 0.906±0.005 | 0.685±0.007 |
| ESC | **0.995**±0.002 | 0.973±0.002 | 0.567±0.002 | 0.881±0.009 | 0.589±0.013 |
| HCT116 | **0.978**±0.004 | 0.931±0.008 | 0.771±0.002 | 0.894±0.007 | 0.586±0.011 |
| HEK293 | **0.980**±0.001 | 0.959±0.004 | 0.854±0.001 | 0.909±0.005 | 0.882±0.005 |
| HEK293T | **0.981**±0.001 | 0.960±0.001 | 0.699±0.001 | 0.889±0.003 | 0.563±0.004 |
| Hela | **0.978**±0.002 | 0.956±0.003 | 0.796±0.002 | 0.895±0.004 | 0.606±0.007 |
| HepG2 | **0.974**±0.002 | 0.950±0.002 | 0.638±0.002 | 0.896±0.005 | 0.518±0.009 |
| MOLM13 | **0.985**±0.001 | 0.944±0.002 | 0.919±0.001 | 0.929±0.003 | 0.937±0.002 |
| Avg. ± Std. | **0.981**±0.002 | 0.953±0.004 | 0.776±0.001 | 0.902±0.005 | 0.698±0.007 |

| **F1** | 3UTRBERT | DeepM6ASeq | WHISTLE | iMRM | SCRAMP |
|---|---|---|---|---|---|
| A549 | **0.963**±0.004 | 0.905±0.008 | 0.860±0.002 | 0.863±0.003 | 0.851±0.005 |
| CD8T | **0.965**±0.003 | 0.947±0.003 | 0.838±0.001 | 0.839±0.006 | 0.580±0.023 |
| ESC | **0.991**±0.002 | 0.960±0.005 | 0.237±0.004 | 0.790±0.026 | 0.626±0.017 |
| HCT116 | **0.969**±0.004 | 0.907±0.008 | 0.703±0.002 | 0.821±0.009 | 0.478±0.054 |
| HEK293 | **0.965**±0.002 | 0.941±0.006 | 0.827±0.001 | 0.848±0.005 | 0.806±0.006 |
| HEK293T | **0.964**±0.001 | 0.956±0.002 | 0.572±0.001 | 0.808±0.004 | 0.404±0.016 |
| Hela | **0.965**±0.003 | 0.946±0.003 | 0.744±0.002 | 0.832±0.004 | 0.540±0.024 |
| HepG2 | **0.965**±0.002 | 0.945±0.002 | 0.438±0.001 | 0.817±0.006 | 0.348±0.076 |
| MOLM13 | **0.965**±0.001 | 0.945±0.002 | 0.910±0.001 | 0.882±0.004 | 0.873±0.003 |
| Avg. ± Std. | **0.968**±0.002 | 0.939±0.004 | 0.681±0.002 | 0.833±0.007 | 0.612±0.025 |

| **MCC** | 3UTRBERT | DeepM6ASeq | WHISTLE | iMRM | SCRAMP |
|---|---|---|---|---|---|
| A549 | **0.928**±0.007 | 0.849±0.017 | 0.776±0.005 | 0.751±0.008 | 0.728±0.009 |
| CD8T | **0.931**±0.005 | 0.913±0.005 | 0.747±0.002 | 0.712±0.010 | 0.316±0.014 |
| ESC | **0.983**±0.004 | 0.941±0.009 | 0.257±0.008 | 0.639±0.016 | 0.172±0.022 |
| HCT116 | **0.939**±0.008 | 0.833±0.017 | 0.602±0.006 | 0.692±0.017 | 0.169±0.024 |
| HEK293 | **0.932**±0.004 | 0.911±0.013 | 0.735±0.003 | 0.725±0.008 | 0.648±0.008 |
| HEK293T | **0.929**±0.001 | 0.912±0.003 | 0.490±0.004 | 0.668±0.007 | 0.126±0.007 |
| Hela | **0.933**±0.005 | 0.913±0.006 | 0.642±0.005 | 0.701±0.007 | 0.183±0.015 |
| HepG2 | **0.932**±0.004 | 0.911±0.004 | 0.388±0.007 | 0.684±0.011 | 0.075±0.010 |
| MOLM13 | **0.931**±0.003 | 0.909±0.005 | 0.843±0.002 | 0.776±0.007 | 0.757±0.007 |
| Avg. ± Std. | **0.938**±0.004 | 0.899±0.009 | 0.609±0.005 | 0.705±0.010 | 0.353±0.013 |

| **ACC** | 3UTRBERT | DeepM6ASeq | WHISTLE | iMRM | SCRAMP |
|---|---|---|---|---|---|
| A549 | **0.963**±0.004 | 0.913±0.008 | 0.877±0.002 | 0.873±0.004 | 0.862±0.005 |
| CD8T | **0.965**±0.003 | 0.946±0.003 | 0.860±0.001 | 0.852±0.005 | 0.650±0.005 |
| ESC | **0.991**±0.002 | 0.960±0.005 | 0.565±0.002 | 0.808±0.009 | 0.582±0.011 |
| HCT116 | **0.969**±0.004 | 0.904±0.009 | 0.769±0.002 | 0.839±0.008 | 0.577±0.007 |
| HEK293 | **0.965**±0.002 | 0.939±0.007 | 0.852±0.001 | 0.859±0.004 | 0.820±0.004 |
| HEK293T | **0.964**±0.001 | 0.945±0.002 | 0.698±0.001 | 0.827±0.003 | 0.554±0.004 |
| Hela | **0.966**±0.003 | 0.945±0.003 | 0.795±0.002 | 0.846±0.004 | 0.589±0.006 |
| HepG2 | **0.965**±0.002 | 0.944±0.002 | 0.637±0.002 | 0.835±0.006 | 0.531±0.006 |
| MOLM13 | **0.965**±0.001 | 0.954±0.003 | 0.917±0.001 | 0.887±0.004 | 0.878±0.003 |
| Avg. ± Std. | **0.968**±0.002 | 0.939±0.005 | 0.774±0.001 | 0.847±0.005 | 0.671±0.006 |

**Supplementary Table. 4.** The average results of the five folds on benchmark data which included six subcellular localizations in terms of AUROC and AUPRC for 3UTRBERT, DM3Loc, RNATracker, mRNALoc and iLoc-mRNA.

| AUROC | 3UTRBERT | RNA_Tracker | DM3Loc | mRNAloc | iLoc-mRNA |
|---|---|---|---|---|---|
| Nucleus | **0.773**±0.014 | 0.739±0.068 | 0.764±0.013 | 0.607±0.009 | 0.498±0.008 |
| Exosome | **0.734**±0.059 | 0.718±0.080 | 0.728±0.060 | 0.410±0.093 | — |
| Cytosol | **0.743**±0.013 | 0.711±0.033 | 0.737±0.018 | 0.509±0.006 | 0.536±0.007 |
| Ribosome | **0.761**±0.005 | 0.730±0.063 | 0.741±0.010 | — | 0.393±0.007 |
| Membrane | **0.756**±0.013 | 0.730±0.041 | 0.753±0.009 | — | — |
| Endoplasmic reticulum | **0.706**±0.012 | 0.570±0.020 | 0.652±0.022 | 0.590±0.016 | 0.421±0.005 |
| AUPRC | 3UTRBERT | RNA_Tracker | DM3Loc | mRNAloc | iLoc-mRNA |
| Nucleus | **0.876**±0.010 | 0.841±0.039 | 0.871±0.009 | 0.765±0.010 | 0.703±0.005 |
| Exosome | **0.997**±0.001 | 0.996±0.002 | 0.996±0.001 | 0.989±0.003 | — |
| Cytosol | **0.323**±0.018 | 0.277±0.048 | 0.322±0.025 | 0.177±0.005 | 0.136±0.003 |
| Ribosome | **0.555**±0.011 | 0.523±0.060 | 0.537±0.014 | — | 0.238±0.002 |
| Membrane | **0.446**±0.021 | 0.379±0.051 | 0.440±0.021 | — | — |
| Endoplasmic reticulum | **0.263**±0.010 | 0.141±0.009 | 0.197±0.018 | 0.142±0.008 | 0.093±0.001 |

**Supplementary Table. 5.** The comparison results on the independent test set which included six subcellular localizations in terms of AUROC and AUPRC for 3UTRBERT, DM3Loc, RNATracker, mRNALoc and iLoc-mRNA.

| AUROC | 3UTRBERT | RNA_Tracker | DM3Loc | mRNAloc | iLoc-mRNA |
|---|---|---|---|---|---|
| Nucleus | **0.726** | 0.662 | 0.716 | 0.626 | 0.526 |
| Exosome | **0.725** | 0.583 | 0.709 | 0.456 | — |
| Cytosol | 0.742 | 0.665 | **0.752** | 0.562 | 0.579 |
| Ribosome | **0.735** | 0.611 | 0.716 | — | 0.421 |
| Membrane | **0.765** | 0.654 | 0.751 | — | — |
| Endoplasmic reticulum | **0.729** | 0.546 | 0.689 | 0.589 | 0.496 |
| AUPRC | 3UTRBERT | RNA_Tracker | DM3Loc | mRNAloc | iLoc-mRNA |
| Nucleus | **0.863** | 0.806 | 0.854 | 0.782 | 0.714 |
| Exosome | **0.992** | 0.985 | 0.991 | 0.981 | — |
| Cytosol | **0.665** | 0.378 | 0.494 | 0.241 | 0.222 |
| Ribosome | **0.563** | 0.427 | 0.537 | — | 0.266 |
| Membrane | **0.524** | 0.321 | 0.505 | — | — |
| Endoplasmic reticulum | **0.351** | 0.154 | 0.294 | 0.179 | 0.131 |
